# Generation of spermatogonia from pluripotent stem cells in humans and non-human primates

**DOI:** 10.1101/2024.05.03.592203

**Authors:** Eoin C. Whelan, Young Sun Hwang, Yasunari Seita, Ryo Yokomizo, N. Adrian Leu, Keren Cheng, Kotaro Sasaki

**Affiliations:** Department of Biomedical Sciences, University of Pennsylvania School of Veterinary Medicine, 3800 Spruce Street, Philadelphia, PA 19104, USA; Department of Pathology and Laboratory Medicine, University of Pennsylvania Perelman School of Medicine, 3400 Spruce Street, Philadelphia, PA 19104, USA; Institute for Regenerative Medicine, University of Pennsylvania, 3400 Civic Center Blvd. Philadelphia, PA 19104, USA

## Abstract

Failures in germline development drive male infertility, but the lack of model systems that recapitulate human spermatogenesis hampers therapeutic development. Here, we have developed a system to differentiate human induced pluripotent stem cells (iPSCs) into primordial germ cell-like cells that self-organize within xenogeneic reconstituted testes (xrTestes) generated from mouse fetal testicular cells. Subsequent transplant of xrTestes into immunodeficient mice resulted in efficient generation of undifferentiated and differentiated spermatogonia as well as preleptotene spermatocytes with striking similarities to their *in vivo* counterparts. As future clinical application will require testing in non-human primates, we utilized a similar strategy to differentiate rhesus iPSCs through all fetal germ cell stages into spermatogonia-like cells. Together, these models will serve as steppingstone to completion of human male in vitro gametogenesis.

## Introduction

Perpetuation of the human species hinges on successful germline development to transmit genetic information to the next generation. Human male germ cells undergo a cascade of developmental steps throughout pre- and postnatal life that culminates in the formation of haploid spermatozoa. While rodent models have been historically informative about the process of male germ cell differentiation, there are many morphological and molecular differences between rodent and primate spermatogenesis (*1*). As errors in germline development can lead to infertility or congenital abnormalities in offspring, understanding the molecular underpinnings of this process in systems that recapitulate human male gametogenesis is imperative.

Recent studies using single cell RNA-sequencing (scRNA-seq) analyses of *in vivo* samples has provided detailed insights into male gametogenesis in humans and non-human primates (*2–11*). During this process, sexually undifferentiated primordial germ cells (PGCs) are specified from pluripotent cells and their subsequent colonization of fetal testes initiates male-specific development, resulting in their differentiation into multiplying (M) prospermatogonia (M prospg) followed by mitotically-arrested primary transitional (T1) prospermatogonia (T1 prospg) (*1*, *9*). Subsequently, T1 prospg transition into two discrete phases of undifferentiated spermatogonia (Spg) (i.e., State 0 [S0] and State 1 [S1] Spg) that likely serve as a founder population for life-long spermatogenesis (*5*, *8*). Although these cells remain largely dormant in pre-pubertal males, onset of puberty triggers differentiation into differentiating Spg (Diff.spg), which subsequently initiate meiosis as spermatocytes (Spc) that give rise to haploid spermatids (*1*).

While scRNA-seq studies of human tissue has provided a wealth of information on the trajectory of male germline development, the mechanisms underlying these changes remain to be evaluated. To this end, we first established a robust directed differentiation method to induce human iPSCs into primordial germ cell-like cells (PGCLCs), which closely resemble pre-migratory stage primate PGCs *in vivo* (*12*). Moreover, by leveraging the self-organizing capacity of testicular somatic cells (*13*), we have integrated 9A13 AGVTPC iPSC-derived PGCLCs into dissociated mouse fetal testicular somatic cells to reconstitute PGCLC-containing 3D testicular architectures termed xenogeneic reconstituted testes (xrTestes) that can be maintained long-term in an air-liquid interface (ALI) culture system (*9*). While PGCLCs matured into cells closely resembling M prospg and T1 prospg within this niche environment, the induction efficiency was low and the testicular tissue organization was compromised after 80 days of culture, likely due to the lack of a blood supply. Accordingly, germ cells beyond T1 prospg stage were not attainable (*9*). To overcome this issue, here we transplanted xrTestes into immunodeficient mice, thereby enabling more sustainable testicular structural reconstitution and derivation of postnatal stage male germ cells up to pre-leptotene spermatocyte. The striking similarities of developing cells to their *in vivo* counterparts provides a valuable model in which to assess mechanisms driving germ line development up to the pre-leptotene spermatocyte stage.

As functional validation of germ cells is limited in humans, producing a non-human primate model is of paramount importance. As rhesus macaques have been established as a vital model for human spermatogenesis, we also established the first iPSC-derived macaque spermatogonia. This system provides us with a parallel population to humans with which to investigate male germ cell development, providing a solid foundation for both preclinical and clinical translational paths towards a regenerative medicine approach for human infertility.

## Results

### Engraftment of xrTestes into NCG mice

Using human iPSCs bearing *TFAP2C/AP2γ-EGFP (AG); DDX4/hVASA-tdTomato (VT); PIWIL4-ECFP (PC)* reporter alleles (9A13 AGVTPC iPSCs), we first induced AG^+^ PGCLCs through iMeLCs (Fig. 1A-C, S1A) (*9*). These cells were reaggregated with mouse fetal testicular somatic cells to generate xrTestes (Fig. S1B). To improve the survival of xrTestes and germ cell maturation, we transplanted xrTestes under the kidney capsule of immunodeficient NCG mice. Remarkably, grafts displayed tubular reconstitution and vascularization one month after transplantation (Fig. S1C). Histological sections revealed well-developed SOX9^+^ seminiferous tubules/cords colonized by EGFP^+^NANOG^+^POU5F1^+^ PGCLCs and interstitial spaces bearing clusters of HSD3B^+^ Leydig cells and CD31^+^ blood vessels (Fig. S1D, E). Reconstituted testes without PGCLC integration revealed empty tubules lacking germ cells, confirming successful depletion of endogenous mouse PGCs (Fig. S1D). Together, these findings indicate the formation of structures inside xrTestes analogous to mammalian seminiferous tubules.

**Figure 1.**
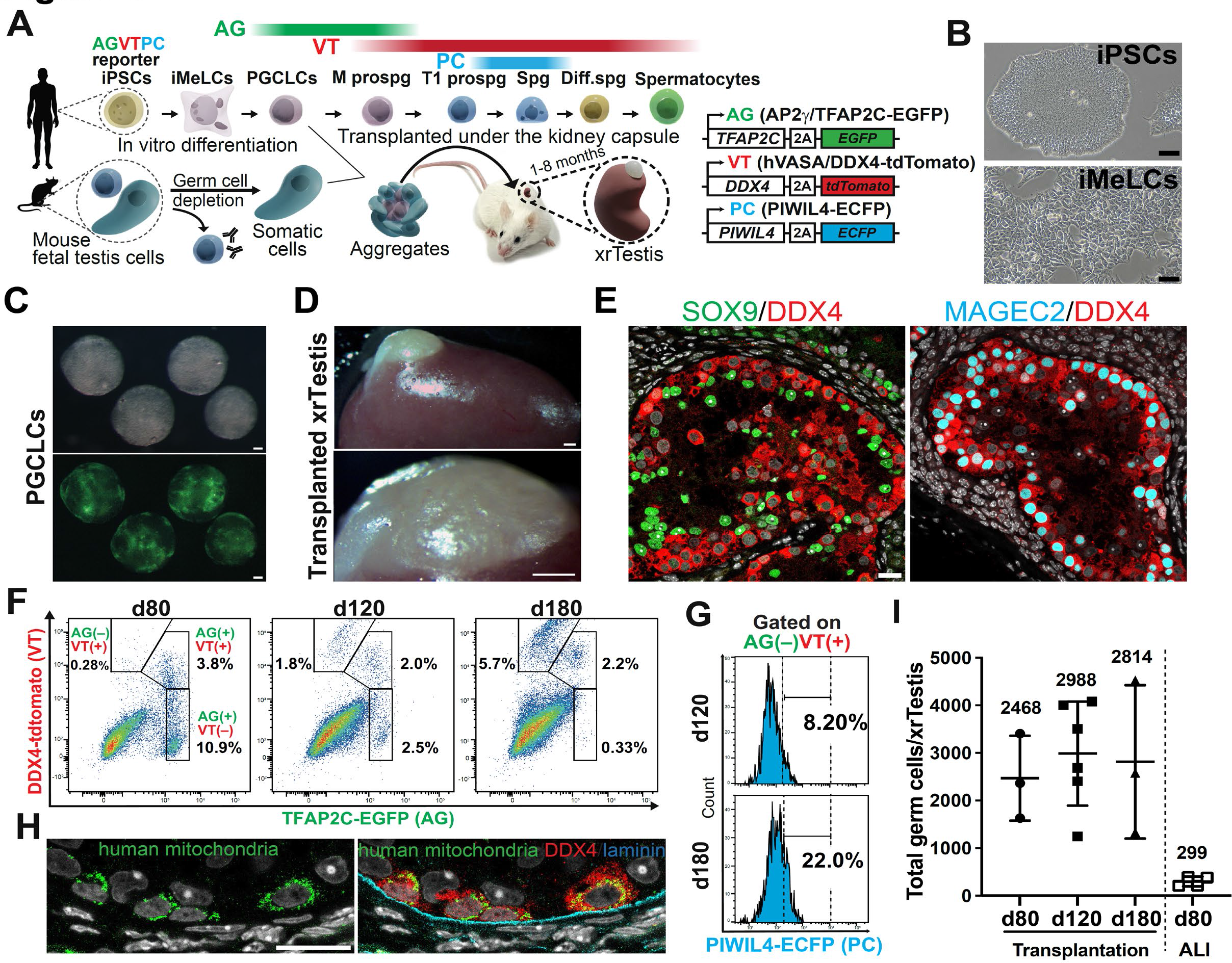
Generation of xenogeneic reconstituted testis (xrTestis) (A) Schematic of experimental design. iPSCs bearing indicated fluorescence reporter alleles were differentiated *in vitro* into first iMeLCs then PGCLCs via a defined culture system. PGCLCs were mixed with mouse fetal testicular somatic cells, aggregated and transplanted under the kidney capsule of NCG mice, forming a xenogenic reconstituted testis (xrTestis) to promote germ cell maturation. (B) Phase contrast images of iPSCs (top) and iMeLCs (bottom) in culture. Scale bar, 50 μm. (C) Bright field (top) or fluorescence images (bottom) of d5 PGCLCs. Scale bar, 200 μm. (D) xrTestis formed on the mouse kidney after six months of transplantation. Scale bars, 500 μm. (E) Immunofluorescence (IF) images of xrTestes for indicated proteins merged with DAPI staining (white). SOX9 marks mouse Sertoli cells. DDX4 and MAGEC2 label cells progressed beyond M prospg and T1 prospg stages, respectively. Scale bar, 20 μm. (F) FACS plots of xrTestes at day(d)80 (left), d120 (middle) and d180 (right) of transplantation. Gates denote AG(+)VT(–), AG(+)VT(+), AG(–)VT(+) populations. (G) Histogram showing proportion of PIWIL4-EGFP (+) cells among AG(–)VT(+) population at d120 (top) and d180 (bottom) of transplantation. (H) IF images of a xrTestis section for human mitochondrial antigen (green), DDX4 (red) and laminin (cyan) merged with DAPI staining (white). Scale bar, 20 μm. (I) The number of total germ cells recovered per xrTestis at indicated time of transplantation as well as generated by ALI culture. Means ± standard deviation.

### Progressive male germ cell maturation in xrTestis

Based on our success, we next carried out longer term xrTestes grafts and found that structural organization of the seminiferous tubules and surface vasculature in the transplanted grafts were still observed 6 months after transplantation (Fig.1D, E). Immunofluorescence (IF) and flow cytometry confirmed the progressive germ cell maturation from AG^+^VT^−^ to AG^+^VT^+^ and AG^−^VT^+^ population, which correspond roughly to *in vivo* PGCs, M prospg and int. prospg/T1 prospg/Spg and their derivatives, respectively (Fig. 1F). Accordingly, a fraction of DDX4/VT^+^ germ cells expressed MAGEC2 or PIWIL4-ECFP (PC), markers of T1 prospg/Spg (Fig. 1E, G) (*9*). Importantly, all DDX4^+^ cells strongly expressed human mitochondrial antigens, further confirming the human origin (Fig. 1H). Similar germ cell maturation was seen when utilizing PGCLCs arising in 2D expansion culture or PGCLCs from a different AGVT reporter iPSC line (18–2) for xrTestis formation (Fig. S2A-G, S3A-I). Consistent with structural preservation of xrTestes, the yield of germ cells per xrTestis was markedly improved in xrTestis transplant compared to ALI culture (∼3000 vs ∼300, ∼10 fold increase) (Fig. 1I).

### Reconstitution of fetal, prepubertal and peripubertal human male germline development *in vitro*

We next sought to elucidate the lineage trajectory and gene expression dynamics of xrTestis-derived germ cells. To this end, we performed scRNA-seq on fluorescently activated cell sorted (FACS) germ cells (AG^+^ or VT^+^) in xrTestes at 4, 6 and 8 months of transplantation and analyzed them together with day 5 PGCLCs (*9*). After filtering out contaminating mouse somatic cells (Fig. S4A), 10,298 cells remained for downstream analysis. Dimension reduction, clustering and projection of these cells onto UMAP revealed 17 clusters, which were annotated based on their differentially expressed genes (DEGs) and previously identified markers unique to each cell type (Fig. 2A-C, S4B, Table S1) (*4*, *5*, *9*, *10*). Accordingly, we identified *POU5F1*^+^ PGCLCs, here defined as PGCs-Early (PGCs.E), and in progression, *NANOS3*^+^ PGCs Late-1 (PGCs.L1), *MLEC1*^+^ PGCs Late-2 (PGCs.L2), *ASB9*^+^ Intermediate (Int.) prospg, *MORC1*^+^*RHOXF1*^+^ T1 prospg, *PIWIL4*^+^*SOHLH1*^+^ Spg, *STRA8*^+^ Diff.spg-Early (Diff.spg.E), *SOHLH2*^+^*SCML2*^+^ Diff.spg-Late (Diff.spg.L) and *MEIOB*^+^*SYCP1*^+^ Preleptotene Spermatocytes (Prelep.spc), which progressed in this order as revealed by RNA velocity and pseudotime analyses (Fig. 2B, C, S4C-F). Consistent with gene expression dynamics *in vivo*, *DDX4* and *DAZL*, markers of gonadal germ cells (*1*, *6*), were gradually turned on as PGCLCs progressed into M prospg (Fig. 2B, S4F, S5, Table S2). Germline specifier (e.g., *TFAP2C*, *PRDM1*, *SOX15*) and pluripotency-associated gene (e.g., *POU5F1*, *NANOG*) expression tapered off as cells progressed into T1 prospg (Fig. S4E, F, S5). Proliferation peaked at PGCs.L/M prospg and then tapered off as cells became Int. prospg/T1 prospg (Fig. 2D). In addition, T1 prospg revealed upregulation of a number of spermatogonial marker genes, some encoding transcription factors (e.g., *SIX1*, *TCF3*) with DEGs enriched for GO terms such as “spermatogenesis” (Fig. 2C, S5). T1 prospg to Spg transition was characterized by further upregulation of spermatogonial marker genes (e.g., *UTF1*, *PIWIL4*, *TCF3*, *SIX1*) whereas cells remained largely quiescent during the transition (Fig. 2B, D, S5). The Spg stage was followed by initiation of differentiation (Diff.spg.E, Diff.spg.L) during which cells resumed proliferation, downregulated spermatogonial markers and modestly upregulated some meiosis-related genes (e.g., *STRA8*, *REC8*) (Fig. 2A-D, S5). Finally, cells transitioned into Prelep.spc by upregulating genes related to meiosis (Fig. 2C, S5). We confirmed the similar lineage progression in xrTestes utilizing either PGCLCs expanded in 2D expansion culture or PGCLCs derived from another iPSC line (18-2 AGVT iPSCs), further supporting the robustness of the platform (Fig. S4G, H).

**Figure 2.**
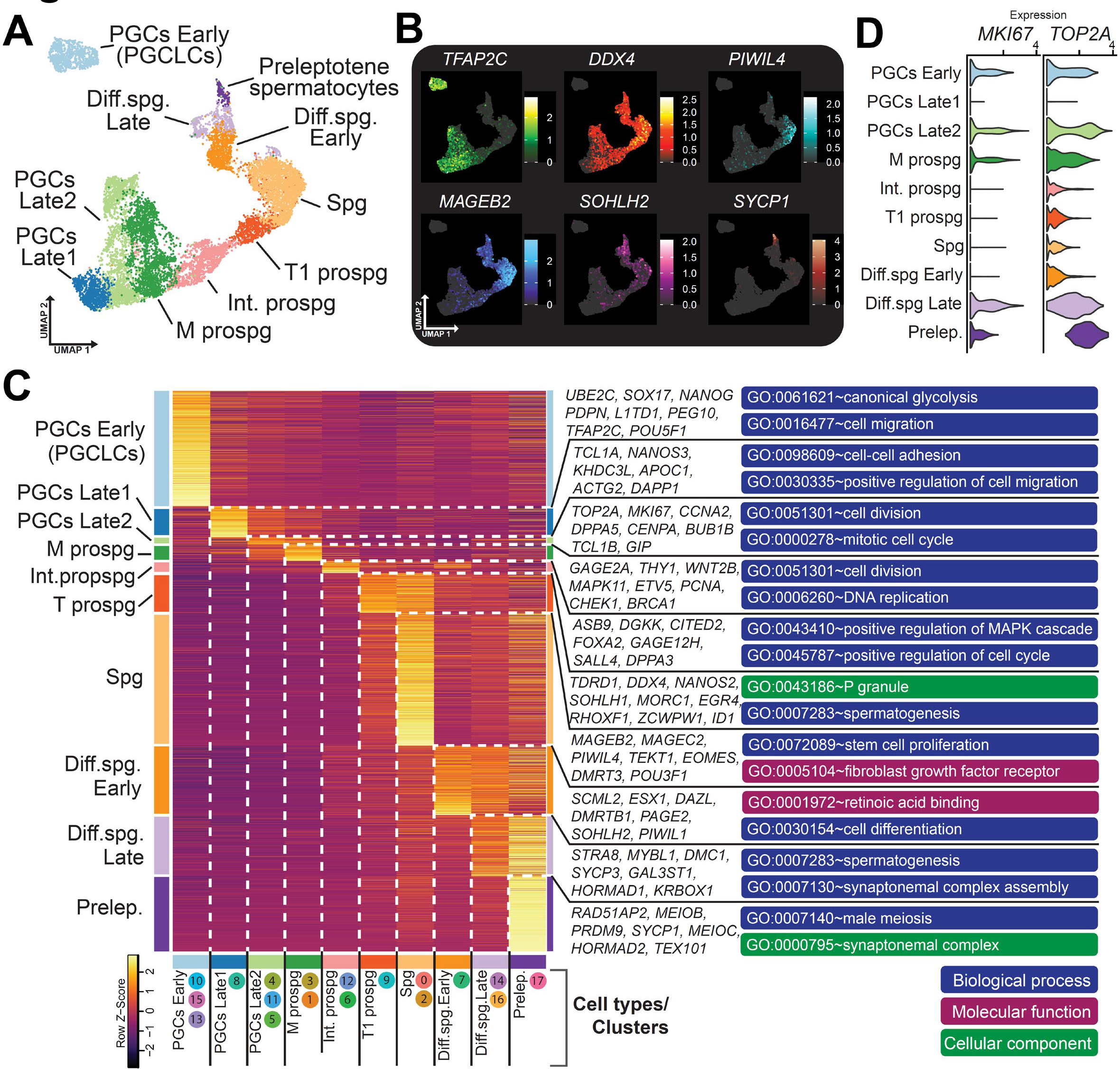
Maturation of iPSC-derived germ cells up to preleptotene spermatocytes within the reconstituted niche. (A) UMAP projection of xrTestis germ cells, with cell types assigned to clusters based upon marker gene expression. (B) Expression of key markers of germ cell differentiation projected on UMAP plot as in (A) (C) Heatmap of the average gene expression for top DEGs for each cell type (unbiased cluster assignments shown below). Select key markers are shown at right along with select GO terms for each set of DEGs, colored by category. (D) Violin plots showing expression of proliferation markers by cell type.

Genome-wide erasure of DNA methylation followed by the acquisition of the androgenic methylation pattern is a hallmark of male gametogenesis. Consistently, we found dramatic genome-wide DNA demethylation as PGCLCs (∼70%) progressed into AG^+^ cells (∼26%), AG^+^VT^+^ (∼18%) and AG^−^VT^+^ (∼12%). Of note, despite the emergence of postnatal type germ cells (e.g., Spg, Diff.spg), AG^−^VT^+^ cells at 6 months of transplantation remained undermethylated (11%), suggesting that *de novo* methylation had not yet commenced in these cells (Fig. S6).

### NANOS3 and DND1 safeguard germline identity during early germline development

Our transcriptomic comparison between PGCs.E and PGCs.L1 revealed upregulation of the RNA-binding proteins *NANOS3* and *DND1,* indicating their potential role in migratory-stage human PGC development (Fig. S7A, B). These genes were also progressively upregulated upon integration of PGCLCs in xrTestis in ALI culture (Fig. S7C, D). To understand the functional significance of *NANOS3*, we generated *NANOS3* biallelic mutant iPSCs from parental 9A13 iPSCs (Fig. S7C). Both mutant and isogenic wild-type iPSCs were readily induced into PGCLCs without overt differences in induction efficiency (Fig. S7E). However, upon xrTestis culture, the number of mutant cells gradually decreased and, by day 14 of ALI culture, most mutant cells disappeared from the tubules (Fig. S7F-I). Overexpression of *NANOS3* in a mutant line during ALI culture rescued germ cell loss, suggesting the critical role of *NANOS3* in germ cell maintenance post-specification (Fig. S7J). Likewise, marked loss of PGCLCs were observed in both *NANOS3*-mutant and *DND1*-mutant xrTestes transplants (Fig. S7K-O). In xrTestes containing *NANOS3*-mutant PGCLCs, there were foci of cells labeled by human mitochondrial antigens but lacking AG labeling, specifically supporting the loss of germ cell identity (Fig. S7P, Q). Remarkably, transcriptome comparison of wild-type and *NANOG* or *DND1* mutant AG^+^ cells revealed upregulation of a number of genes related to neuronal differentiation in mutant lines (e.g., *ASCL2, PCP4, SOX11, TUBB2B, NECAB1, LIN28A*) (Fig. S7R, S). These findings suggest that *NANOS3* and *DND1* safeguard germ cell identity by suppressing neuronal dedifferentiation and highlight the capacity of our xrTestes system to provide crucial mechanistic insight into male gametogenesis.

### Validation of xrTestis-derived male germ cells

To further validate the iPSC-derived germ cell progression captured by scRNA-seq, we performed IF combining different markers. We noted that a small number of AG^+^NANOG^+^VT^−^ cells persisted at 4 months, consistent with residual PGCLCs (Fig. 3A, B). In 6 months xrTestes, we identified TFAP2C^+^DDX4^+^ cells expressing KIT^+^, in line with PGCs.E/PGCs.L/M prospg (Fig. 3A-B). Consistent with numerous ISH signals for *PIWIL4* (a marker of T1 prospg/Spg) within the reconstituted tubules, many UTF1^high+^TFAP2C^−^ Spg were also identified by IF (Fig.3A-C). We also identified KIT^+^DDX4^low+^MKI67^+^ Diff. spg along the basement membranes that could be distinguished from PGCs/M prospg by the lack of TFAP2C (Fig. 3A, B, D, S8). Furthermore, we identified rare MAGEA3^+^DDX4^+^ cells expressing SYCP3, REC8 and γH2AX by IF and *MEIOB*^+^ cells by ISH, suggesting the emergence of Prelep.spc (Fig. 3A, B, E, S8). The finding is consistent with scRNA-seq data showing the presence of *MEIOB^+^* Prelep.spc at the end of the trajectory (Fig. 3F-G). Additionally, these transcripts were confirmed to be of human origin, providing further support for the successful derivation of Prelep.spc from human iPSCs (Fig. 3H).

**Figure 3:**
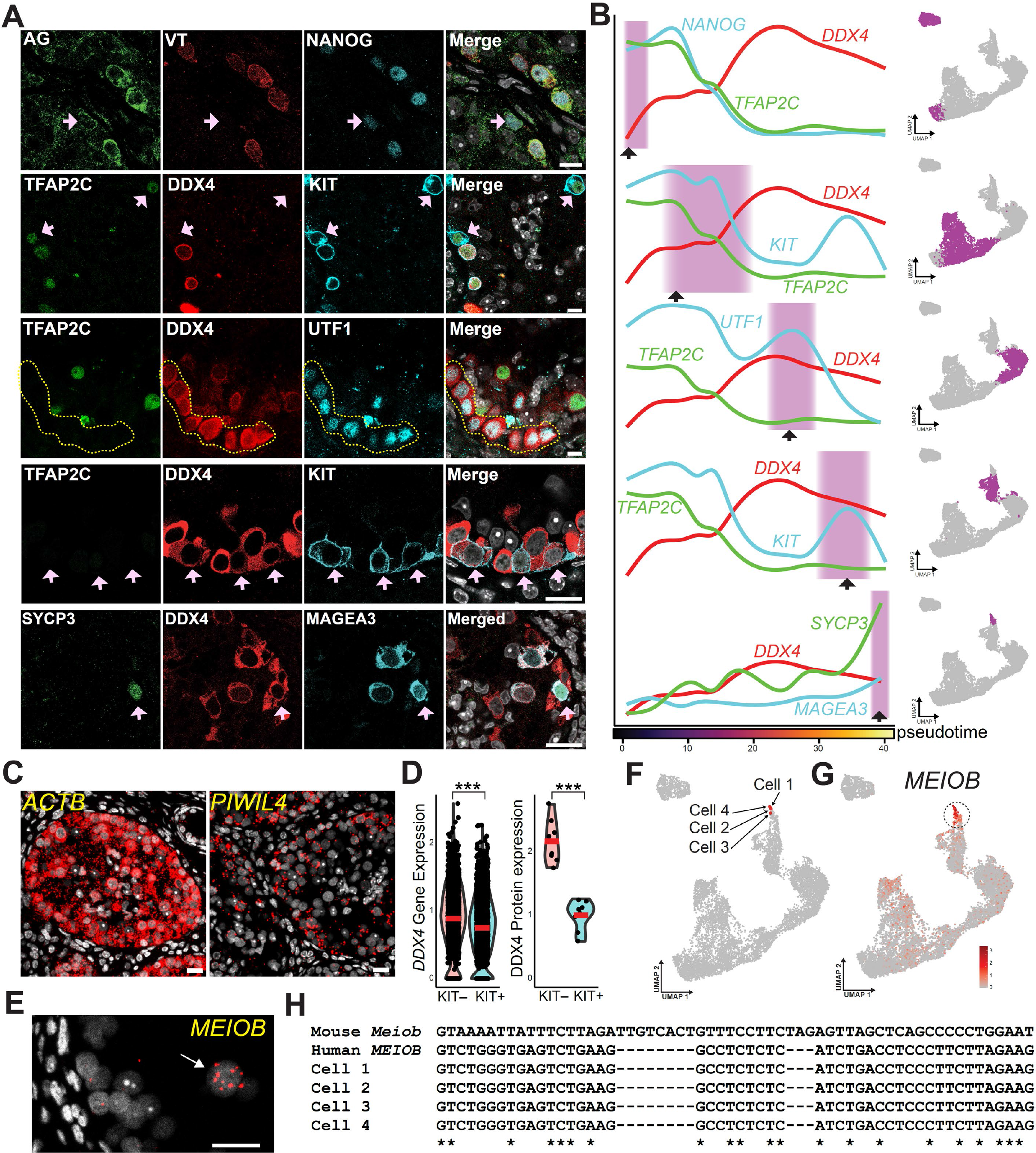
Progressive maturation of germ cells within xrTestis validated by IF and in situ hybridization. (A) IF of xrTestis sections for key markers corresponding to distinct differentiation stages of germ cells: from top to bottom, PGCs (pink arrows, scale bar 10 μm, 4 months), PGCs/M prospermatogonia (pink arrows, scale bar 10 μm, 4 months), undifferentiated spermatogonia (yellow dotted lines, scale bar 10 μm, 6 months), differentiating spermatogonia (pink arrows, scale bar 20 μm, 6 months) and preleptotene spermatocytes (pink arrows, scale bar 20 μm, 4 months). (B) Average gene expression of genes corresponding to IHC markers in (A) along pseudotime. Black arrows indicate approximate in match in pseudotime to pink arrows in (A). Shaded areas correspond to highlighted clusters in UMAPs (right). (C) ISH of *ACTB* (housekeeping gene) and *PIWIL4*. (D) *DDX4* gene expression in *KIT^+^* and *KIT^−^* TFAP2C*^−^* cells as determined by single-cell RNAseq (left). Relative intensity of DDX4 fluorescence via IF of cells scored manually as KIT^+^TFAP2C*^−^* and KIT^+^TFAP2C*^−^*. *** *p* < 0.001, Student’s *t-*test, bars indicate mean. (E) ISH of *MEIOB* transcripts detected in rare cells located towards the lumen of tubules. (F) Location of four representative cells containing *MEIOB* transcripts with 100% match to the human *MEIOB* gene. (G) Expression of *MEIOB* in all xrTestis cells. (H) Alignment of 3’ portions of *MEIOB* transcripts from the four representative cells shown in (F) with human and murine references.

### Human iPSC-derived male germ cells in xrTestis recapitulate normal developmental trajectory in vivo

To evaluate the normalcy of xrTestis-derived germ cells, we integrated the transcriptomes of xrTestis-derived germ cells and our atlas of *in vivo* human germ cell development from migrating PGCs to round spermatids (*10*). This analysis revealed that xrTestis-derived germ cells and in vivo germ cells clustered together based on cell types rather than sample origin and formed a uniform trajectory (Fig. 4A, B, S9A). This analysis also confirmed that the trajectory of xrTestis-derived germ cells was arrested at the Prelep.spc stage. Stage-by-stage comparison demonstrated high concordance between *in vivo* and xrTestis-derived cell types, among which Spg had the lowest number of DEGs and the highest correlation (R2 =0.97, Fig. 4C-E, S9B, C, S10, Table S3). Accordingly, pathway enrichment analysis by cell type revealed similar enrichment patterns between *in vivo* and xrTestis. For example, Spg both *in vivo* and in xrTestis exhibited enrichment of pathway terms such as “ERK/MAPK signaling” and “GDNF Family Ligand-Receptor Interactions”, key signaling pathways in rodent Spg maintenance (Fig. S9C) (*1*, *14*). Notably, further sub-clustering of Spg revealed two previously identified subtypes, *PIWIL4*^+^*MSL3*^+^ State 0 (S0) and *ETV5*^+^*L1TD1*^+^ State 1 (S1) spg in both xrTestis and in vivo testes (Fig. S11A-E) in which the striking transcriptional similarities between xrTestis- and *in vivo*-derived cells highlighted the precise recapitulation of human male gametogenesis in xrTestis (Fig. S11E). Pseudotime trajectory analysis revealed that T1 prospg first progressed into S0 spg, which then transitioned into S1 spg followed by Diff.spg.E, suggesting that S0 spg represents the most undifferentiated Spg (Fig. S11A). Pairwise transcriptomic comparison between S0 and S1 spg revealed highly similar sets of DEGs in both *in vivo* and xrTestis (Fig. S11F, G). Notably S1 spg upregulated a number of mitochondrial genes enriched with GO terms such as “mitochondrial inner membrane”, suggesting that a metabolic switch to oxidative phosphorylation, the dominant pathway in Diff.spg, might already be initiated at S1 spg (Fig. S11 F, H).

**Figure 4.**
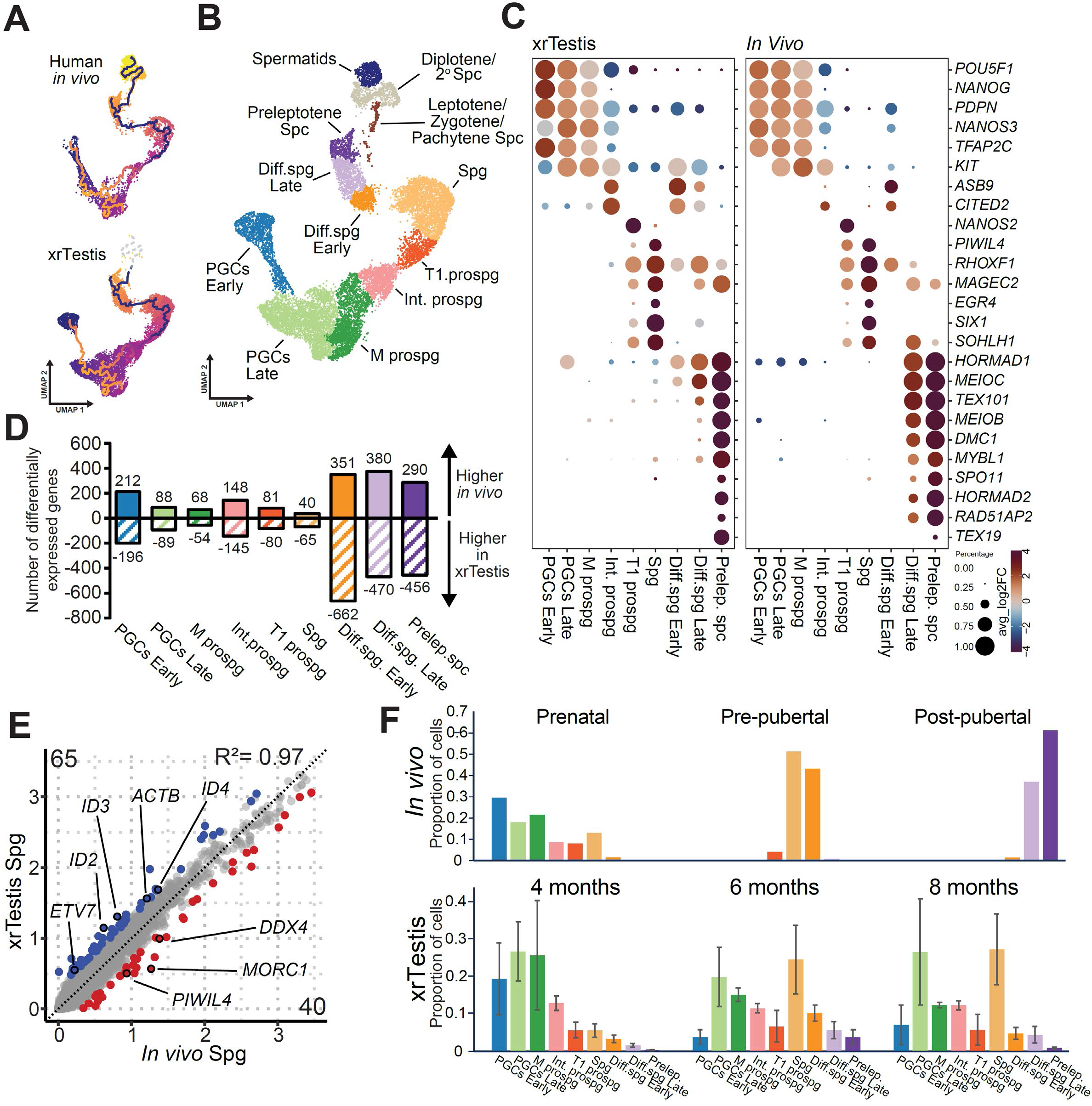
xrTestis germ cells showed differentiation similar to *in vivo* human germ cells. (A) UMAP plots showing human *in vivo* and xrTestis-derived germ cells integrated together and pseudotime calculated. (B) Cell types assigned to joint cell clustering. (C) Dot plot of key gene expression by cell type. The size of the dot indicates the percentage expression of the gene in question for that cell type and color displays the mean log_2_ fold-change as compared with all other cell types. (D) Number of DEGs (pairwise comparison between in vivo and xrTestis-derived germ cells) by cell type (-0.25 > log_2_ fold change > 0.25, *adjusted p value* < 0.05). (E) Scatter plot comparing the average gene expression between xrTestis spermatogonia vs *in vivo* spermatogonia. Number of genes that are differentially expressed are indicated in the corners of the plot, highlighted in blue or red as appropriate. Select genes of interest are shown. (F) Upper panel: proportion of cells of each type derived from prenatal, pre-pubertal, and post-pubertal human samples. Lower panel: proportion of cells derived from xrTestes transplanted for four, six and eight months respectively, error bars represent standard error of the mean.

The number of DEGs increased and the correlation decreased as Spg became Diff.spg and Prelep.spc (Fig. 4D). These DEGs revealed upregulation of genes related to RA signaling in xrTestis-derived germ cells, suggesting a potential alteration in RA signaling in the xrTestis environment (Fig. S10, Table S3).

4 month-old xrTestes showed a dominance of PGCs and prospermatogonia, broadly resembling the prenatal human testes, while 6 month-old xrTestes demonstrated dominance of Spg, similar to pre-pubertal testes *in vivo*. However, unlike pre-pubertal testes, 6 month-old xrTestes exhibited a marked heterogeneity, containing a wide variety of cell types including PGCs/M prospg and Diff.spg.L/Prelep.spc that are normally only present in prenatal and post-pubertal testes respectively. Such findings are suggestive of a differentiation block at the prospg stage and the presence of pro-differentiation signals at Spg stage to facilitate differentiation up to preleptotene stage in xrTestis microenvironment. As extension of transplantation period to 8 months did not significantly alter the cellular composition nor promote further differentiation (Fig. 5E), our findings suggest that signals promoting further differentiation are lacking in this system.

**Figure 5:**
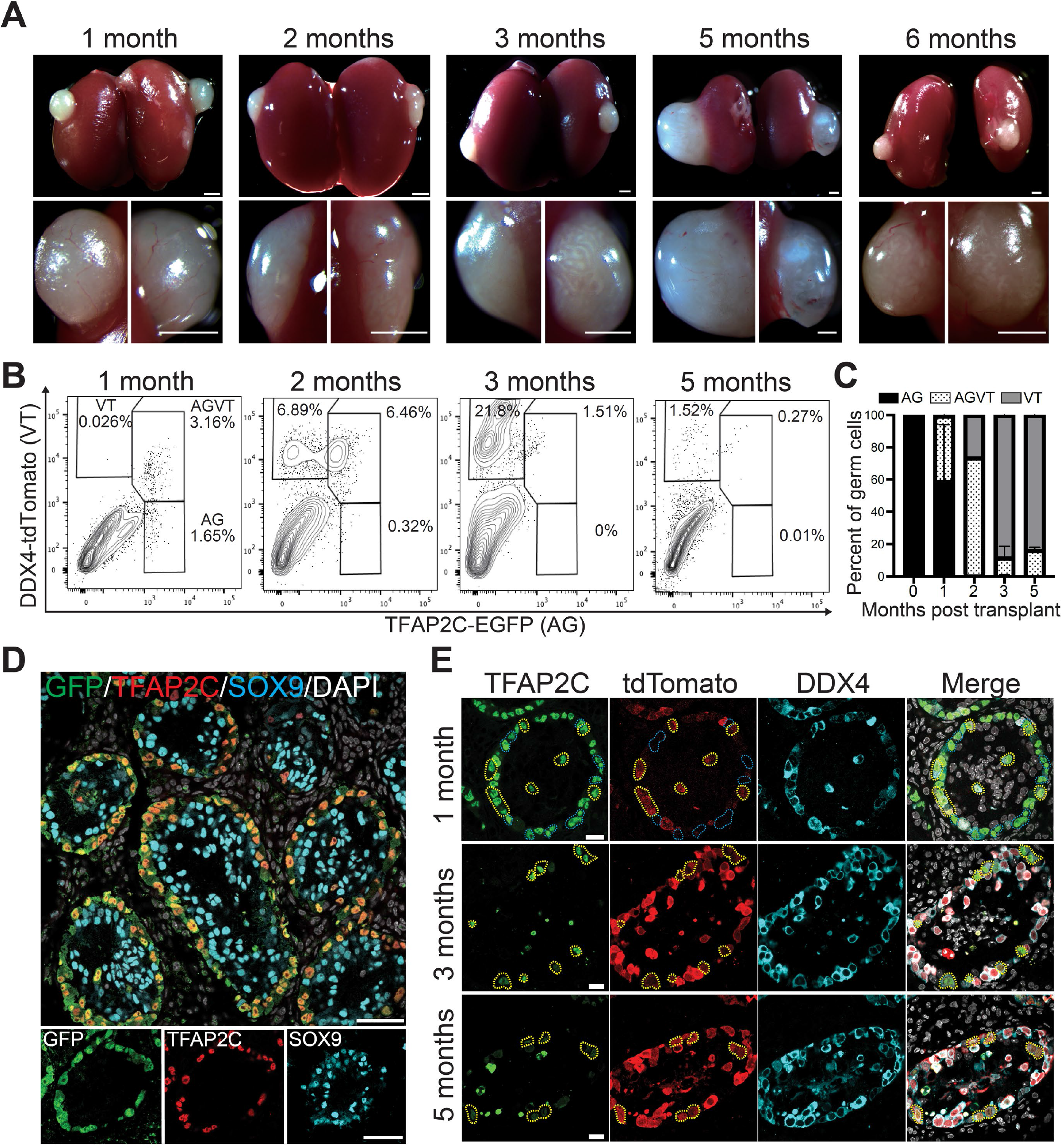
Generation of xrTestes for maturation of rhesus macaque iPSC-derived male germ cells. (A) xrTestes derived from rhesus macaque iPSCs (1C2AGVT19 line) carrying TFAP2C-EGFP (AG) and DDX4-tdTomato (VT) reporter alleles. Scale bars, 1mm. (B) FACS plots of xrTestes at various times post-transplantation. (C) Proportions of AG/AGVT/VT gated fractions in pre-transplantation xrTestes and for each analysis in part (B) (D) IF of 1-month xrTestis for GFP (indicating TFAP2C-EGFP reporter expression, green), TFAP2C (red) and SOX9 (cyan) merged with DAPI staining (white). SOX9 marks mouse Sertoli cells. Scale bars, 50 μm. (E) IF of representative tubules at one, three and five months, detecting TFAP2C protein (green), *DDX4-*tdTomato reporter (red) and DDX4 protein (cyan). Merged images are shown at right. Blue dotted lines indicate TFAP2C+DDX4- cells (i.e. PGCs), whereas yellow dotted lines outline double-positive cells (i.e. M prospermatogonia). Scale bars, 20 μm.

### Reconstitution of male germline development using rhesus macaque iPSCs

While our human iPSC system provided valuable insight into male gametogenesis, future studies of fertility competency must be carried out in non-human primates. Therefore, we next tested whether similar spermatogonial induction could be attained using iPSCs of rhesus macaques in our xrTestis system. Rhesus macaque iPSCs bearing AGVT reporter alleles revealed robust induction into PGCLCs using the 3D culture method described previously (Fig. S12A-I) (*15*). However, because xrTestis generated using AG^+^ PGCLCs developed into teratomas following transplantation into NCG mice (Fig. S12J-L), we performed 2D expansion culture of PGCLCs in the presence of Forskolin, SCF and FGF2 (Fig. S13A-F) (*16*). This culture successfully maintained PGCLCs up to 75 days without altering key pluripotency-associated and PGC markers (Fig. S13G). Post-expansion culture PGCLCs showed higher recovery of germ cells following floating culture and prevented teratoma formation after transplantation (Fig. S12K, L). These xrTestes stably maintained structural organization up to 7 months and exhibited progressive differentiation of iPSC-derived germ cells from the AG^+^VT^−^ to the AG^−^VT^+^ state, suggesting the successful derivation of T1 prospg/Spg (Fig. 5A-E).

We next FACS-isolated AG^+^ or VT^+^ germ cells from xrTestis explants for scRNA-seq analyses. Projection of germ cells onto UMAP after removal of mouse cells revealed progressive germ cell differentiation in which *TFAP2C*^+^*DDX4*^−^ PGCs.E gradually transitioned into *TFAP2C*^−^*DDX4*^+^*PIWIL4*^+^ Spg, similar to human xrTestes (Fig. 6A-F, Table S4). Because *in vivo* transcriptomic references of rhesus macaque during fetal stages have not been well established, we compared the trajectory to the human trajectory we established earlier. This analysis revealed that rhesus iPSC-derived germ cells progressed similarly to both humans in vivo and human xrTestis, albeit spermatocytes were lacking in rhesus xrTestes (Fig. 6G). These findings indicate the conservation of developmental niche signals in testicular somatic cells that allow robust differentiation of Spg from PGCLCs (Fig. 6H). Because rhesus macaque Spg are competent to generate spermatids upon seminiferous tubule transplantation (*17*), our system may also allow validation of the functionality of iPSC-derived wild type and mutant primate male germ cells in future mechanistic studies.

**Figure 6.**
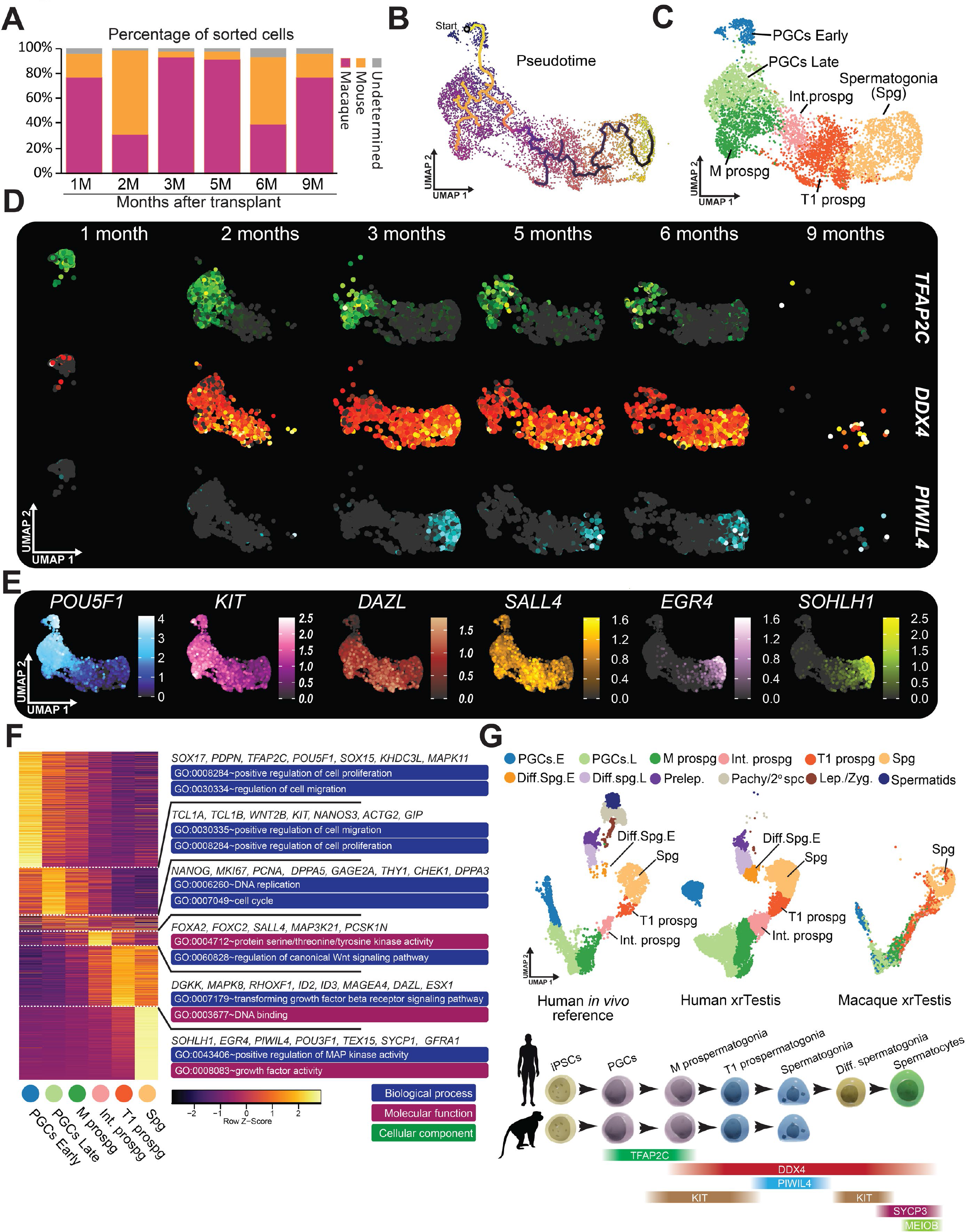
Macaque iPSC-derived germ cells differentiate into spermatogonia (A) Proportion of captured macaque cells relative to contaminating mouse cells as determined by alignment algorithm. (B) Pseudotime progression of iPSC-derived germ cells. (C) Cell type assignments to unbiased clusters based on marker gene expression. (D) Expression of *TFAP2C, DDX4* and *PIWIL4* split by transplantation time. (E) Expression of additional key germ cell markers. (F) Heatmap of top genes expressed by cell type. Select genes are shown for each cell type along with representative GO terms. (G) Label transfer of macaque cells using human *in vivo* and xrTestis cells as a reference. (H) Model of *in vitro* germ cell derivation using human and rhesus macaque i

By reconstituting testicular niche and germ cell interaction in xrTestis, we have successfully induced both human and rhesus macaque iPSCs into Spg, putative stem cells responsible for life-long spermatogenesis through their self-renewal and differentiative capacity. Remarkably, Spg induced from human iPSCs were capable of further differentiating into preleptotene spc, further lending support to the functionality of iPSC-derived Spg. However, differentiation has not progressed beyond the preleptotene stage, despite extension of the transplantation period up to 8 months. This could be due to lack of paracrine or hormonal signaling (e.g., gonadotropin, testosterone) or inefficient Sertoli cells-germ cell interactions required for meiotic progression.

Robust generation of iPSC-derived Spg opens up new possibilities for interrogating the molecular circuitry required for their development using both genetic and pharmacologic approaches. Because rhesus macaque Spg can generate spermatids following seminiferous tubule transplantation (*17*), findings in our platform may also allow validation of the functionality of iPSC-derived primate male germ cells, i.e., generation of fertilization-competent haploid spermatids in future studies. Finally, this platform will provide an unlimited source of genetically-traceable human undifferentiated Spg, allowing us to identify culture conditions suitable for expanding self-renewing Spg in humans. Such cultures will ultimately enable us to understand the molecular basis of stemness in primate Spg and could be used to inform future infertility treatments.

## Acknowledgments

We thank Drs. Leslie King for carefully reviewing the manuscript and providing insightful comments. We thank members of Sasaki lab for the discussion of this study. We acknowledge the Comparative Pathology Core at the University of Pennsylvania School of Veterinary Medicine for the preparation of paraffin sections.

## Funding

Open Philanthropy projects, 10072620 (KS)

Open Philanthropy projects, 10095457 (KS)

Open Philanthropy projects, 10080664 (KS)

The Heath Research Formula Funds, the Pennsylvania Department of Heath 67-80 (KS)

## Author contributions

Conceptualization: KS

Methodology: KS

Investigation: KS, ECW, YH, YS, RY, NAL, KC

Visualization: KS, ECW, RY

Funding acquisition: KS

Project administration: KS

Supervision: KS

Writing – original draft: KS, ECW, RY, YS, YH, KC

Writing – review & editing: KS, ECW

## Competing interests

Authors declare that they have no competing interests.

## Data and materials availability

All data are available in the manuscript or the supplementary materials. Materials and reagents are either commercially available or available upon request from the correspondence author. scRNA-seq data have been deposited in the Gene Expression Omnibus (GEO) database (GSE266123).

## Supplementary Materials

Materials and Methods

Figs. S1 to S13

Table S1 to S6

References (18-25)

## Notes

### Competing Interest Statement

The authors have declared no competing interest.

