## Supplementary Materials for "Generation of spermatogonia from pluripotent stem cells in humans and non-human primates"

**Materials and Methods**

#### Culture of human iPSCs

Human iPSCs were cultured on six-well plates coated with recombinant laminin-511 E8 (iMatrix-511 Silk, Nacalai USA) and were maintained under a feeder-free condition in StemFit® Basic04 medium (Ajinomoto) containing basic FGF (Peprotech) at 37ºC under an atmosphere of 5% $CO_2$ in air. Prior to passaging or the induction of differentiation, iPSC cultures were treated with a 1:1 mixture of TrypLE Select (Life Technologies) and 0.5 mM EDTA/phosphate-buffered saline (PBS) for 15 min at 37ºC to dissociate them into single cells. Subsequently, 10 μM ROCK inhibitor (Y-27632; Tocris) was added in the culture medium for 1 day after passaging iPSCs.

#### Generation of human iPSCs bearing AGVT fluorescence reporter alleles

The TALEN constructs and donor vectors for generating the *TFAP2C/AP2γ-p2A-EGFP* (AG); *DDX4/human VASA-p2A-tdTomato* (VT) alleles were described previously (*1*, *2*). TALEN's RVD sequences were as follows: TFAP2C-left (5-prime), NN NN NI NN NI NI NI HD NI HD NI NN NN NI NI NI NG; TFAP2C-right (3-prime), NI HD NG HD NG HD HD NG NI NI HD HD NG NG NG HD NG. DDX4-left, HD HD HD NI NI NG HD HD NI NN NG NI NN NI NG NN NI NG NN; DDX4-right, NN NI NI NN NN NI NG NN NG NG NG NG NN NN HD NG NG. To establish AGVT knock-in fluorescence reporter human iPSCs, we first transfected the donor (5 ug) and TALEN-expression vectors (2.5 ug each) for VT into one million iPSCs (clone Penn083i-679Sev4 derived from peripheral blood erythroblasts of male donor, kind gift from Dr. Wenli Yang) (*3*) by NEPA21 Type II Electroporator (Nepagene), followed by neomycin selection. Puromycin-resistant colonies were pooled, expanded then subsequently transfected by donor and TALEN-expression vectors targeting AG as described above, followed by puromycin selection and transfection of a plasmid expressing Cre recombinase to remove the PGK-Puro and PGK-Neo cassettes. The success of the targeting with Cre recombination and without random integration events were confirmed by PCR with extracted genomic DNA from each colony. One clone bearing heterozygous AG and homozygous VT alleles (18-2) were further confirmed to be devoid of indel within the non-recombined *TFAP2C* allele. Another AGVT knock-in reporter iPSCs (1719) were generated using 585B1-1-7 (human iPSC line bearing heterozygous AG allele, kind gift from Drs. Sasaki and Saitou at Kyoto University) as parental iPSCs (*1*). We transfected the TALEN constructs and the donor vector for VT and obtained one clone (designated as 1719 iPSCs) bearing heterozygous VT allele without random integration or indel formation in the non-recombined allele by using the same approach as above. Normal karyotypes were confirmed by G-banding analyses as described in elsewhere.

**Generation of *NANOS3* and *DND1* knock-out human iPSCs**

Two pairs of sgRNA sequences before and after the targeted sequence of *DND1* and NANOS3 were designed with CRISPR guide RNA design tool (Benchling). One pair of sgRNAs targeting *DND1* (TATCCCGCTGTTCCAGCGCG and TGGTGCTCGTACACGTCCTG for exon3) or two pair of sgRNAs targeting *NANOS3*

(ACCTGGTTAGGGCTCTGAGT and GTAATCTGTCCACAGGTCAA for pre-exon1, and GCCTGACAAGGCGAAGACAC and GGTGGTGTGGCTGTAGACGG for post-exon1) were selected and cloned into the *pX335-U6-Chimeric BB-CBh-hSpCas9n (D10A) SpCas9n*-expression vector to generate the sgRNAs/Cas9n vector (Addgene, #42335) (*4*). Four sgRNAs/nCas9 vectors (2.5 µg each) were transfected into 1 million AGVT iPSCs (9A13 or 1719) by electroporation with a NEPA21 Type II electroporator (Nepagene). Two days after transfection, mCherry-expressing cells were sorted into a 96-well plate (single cell/well) with BD FACSAria Fusion, and subsequently expanded and genotyped by PCR. The deletions in *DND1* and *NANOS3* were further validated by sequencing.

**Generation of DND1 and NANOS3 expression lines**

The vectors carrying the Doxycycline-induced expression cassette were constructed using the Gateway System (Thermo Fisher Scientific) as described previously (*5*). The full-length cDNA sequences of *DND1* and *NANOS3* were PCR amplified from d5 human PGCLCs derived from the 9A13 AGVT iPSC line. Nucleotide sequences for the epitope tags with a linker, 3×FLAG-G4S were added to the 5' ends of *DND1* and *NANOS3*. The PCR products were cloned between the BamHI and XhoI sites of the pENTR 1A dual selection vector and were subsequently recombined into the destination vector with a LR Clonase II Enzyme mix kit (Thermo Fisher). In the destination vector, the transgenes were cloned under the TetO promoter repeat region, which was followed by the rabbit β-globin poly A sequence. In the region downstream of rabbit β-globin poly A sequence, the puromycin-resistant gene driven by the EF1α promoter was inserted.

Transfection was performed with the electroporator NEPA21 type II (Nepagene). One million iPSCs were transfected with 500 ng PiggyBac transposase expression vector and 1 μg transgene expression vector, then resuspended in 100 μl of OptiMEM (Thermo Fisher Scientific). Selection antibiotics (0.2 μg/ml puromycin) were added two days after the transfection and maintained until the surviving colonies were picked up at d12–14. The induction of the transgenes with 1.0 μg/ml doxycycline (Takara-Clontech) for the selected iPSC clones was assessed at 24 h of culture.

**Cell Collection and reprogramming to rhesus macaque iPSCs**

Rhesus monkey fibroblasts were gifted by Dr. J. Hennebold (the Oregon National Primate Research Center). Rhesus monkey fibroblasts were reprogrammed using the CytoTune-iPS 2.0 Sendai Reprogramming Kit (Thermo Fisher) according to the manufacturer's instructions. Briefly, $4 \times 10^4$ fibroblasts were plated in each well of a six well plate and cultured in Dulbecco's Modified Eagle Medium (DMEM) containing 10% fetal bovine serum (FBS) one day before infection. On the first day (day 0), Sendai viruses containing KOS, C-Myc and KLF-4 expression cassettes were added to the fibroblasts at a 5:5:3 ratio and these cells were cultured until day 7 in DMEM containing 10% FBS. On day 7, cells were plated onto a mouse embryonic fibroblast feeder (MEF) layer ($2.5 \times 10^5$) in DMEM containing 10% FBS. Medium was changed daily until day 8 when the media was switched to AITSIF20 (advancedRPMI1640 and Neurobasal [1:1] supplemented with AllbuMax [1.6%], 1× ITS [Insulin, Transferrin, Selenium], IWR1 [2.5 μM], and bFGF

[20 ng/ml]) (*6*, *7*). Between days 14 and 28, individual colonies became visible, which were handpicked and transferred individually to a new well containing MEF. These were passaged manually for several passages until cryopreservation. After this point they were with passaged using a 10 minute incubation in 0.5 x Triple select in PBS. G-band karyotype analyses were performed using Cell Line Genetics (Madison, WI).

**Culture of rhesus macaque iPSCs**

For a feeder free iPSC culture, iPSCs (1C2) were cultured on a xeno-free recombinant Laminin-511 E8 fragment-coated dish (TAKARA, iMatrix-511silk) with AITSIF20 containing 50 ng/ml Activin A, 3 uM CHIR9921 and 30 uM Y27632 media. The cells were passaged around every 6–7 days with 0.5 x Triple select in PBS. For on feeder culture, iPSCs were cultured with AITSIF20 on MEFs ($2.5\times10^5$cells/well of a 6-well plate) and culture medium was supplemented with 10 μM Y27632 inhibitor until 24 h after passage.

**Generation of rhesus macaque iPSC lines bearing knock-in fluorescent reporter alleles**

To construct the donor vectors for generating rhesus macaque iPSC lines bearing *TFAP2C-P2A-EGFP* and *DDX4-P2A-tdTomato* alleles, the homology arms of *TFAP2C* and *DDX4* were amplified via PCR with the rhesus monkey fibroblast genome as a template and the primer pairs described elsewhere (*6*). The amplified fragments were subcloned into the *pCR2.1* vector using a TOPO TA Cloning kit (Thermo Fisher, K4500-

40). The *p2A-EGFP* or *p2A-tdTomato* fragments with *pgk-Puro* or *pgk-Neo* cassettes flanked by the *loxP* sites, respectively, were amplified by PCR from the targeting vectors used for the generation of 18-2 AGVT human iPSCs. These fragments were inserted in-frame upstream of the stop codons of *TFAP2C* or *DDX4* in the pCR2.1 vector containing the homology arms, using a GeneArt Seamless Cloning & Assembly Kit (Thermo Fisher, A13288).

Guide RNAs for the targeting of the *TFAP2C* and *DDX4* loci were designed, as illustrated in Fig. S12. For each guide RNA, two oligonucleotides with guide sequences and BbsI-compatible ends were annealed and cloned into the BbsI site of the *pX335-U6-Chimeric BB-CBh-hSpCas9n (D10A)* vector.

To introduce both targeting vectors into rhesus macaque iPSC line, $1 \times 10^6$ 1C2 rhesus macaque iPSCs were suspended in 100 μl OptiMEM (Thermo Fisher, 31985062) and first transfected with 1 μg each of the Cas9n vector and 3 μg of the donor vector targeting the *TFAP2C* locus using the NEPA21. These cells were then seeded onto DR4 MEFs (ATCC, SCRC-1045). Two days post-transfection, the cells were cultured in AITSIF20 medium, supplemented with 1.0 μg/ml puromycin (Thermo Fisher, A1113803) for 8–10 days. Subsequently, $1 \times 10^6$ puromycin-resistant cells were transfected with 1 μg each of the Cas9n vector and 3 μg of the donor vector targeting the *DDX4* locus, then seeded onto a dish containing DR4 MEFs. Two days later, the cells were cultured in the presence of 200 μg/ml G418 sulfate (Gibco, 10131035) and selected over a period of 8–10 days. The surviving cells were then transfected with the vector expressing the Cre recombinase and plated onto dish containing wild-type MEFs, and the resultant colonies were picked and expanded for the genotyping and G-banding analysis.

##### Induction of PGCLCs

Human PGCLCs were induced from human iPSCs via iMeLCs as described previously (*1*, *2*). For the induction of iMeLCs, iPSCs were plated at a density of 4–5 × $10^4$ cells/$cm^2$ onto a human fibronectin (Millipore)-coated 12-well plate in GK15 medium (GMEM [Life Technologies]) with 15% KSR, 0.1 mM NEAA, 2 mM L-glutamine, 1 mM sodium pyruvate, 100 U/ml PenStrep, and 0.1 mM 2-mercaptoethanol) containing 50 ng/ml of ACTIVIN A (R&D Systems, 338-AC), 3 μM CHIR99021 (Tocris Bioscience, 4423) and 10 μM of Y-27632 (Tocris, 1254). After 31–38 h, iMeLCs were harvested and dissociated into single cells with TrypLE Select. Cells were then plated into Cell repellent V-bottom 96-well plates (Greiner) at 3500 cells per well for induction into PGCLCs. Wells contained GK15 medium supplemented with 200 ng/ml BMP4 (R&D Systems, 314-BP-010), 100 ng/ml SCF (R&D Systems, 255-SC-010), 50 ng/mL EGF (R&D Systems, 236-EG), 1000 U/ml LIF (Millipore, LIF1010) and 10 μM of Y-27632.

Rhesus macaque PGCLC induction were followed as described previously with minor modification (*6*). Briefly, cells were treated with 0.5× TrypLE Select in PBS to make single-cell suspensions. PGCLCs were induced by plating 3,500 iPSCs into a well of Cell repellent V-bottom 96-well plates, in aRB27 [Advanced RPMI 1640 (Thermo Fisher, 12633-012), 2× B27 (Thermo Fisher, 17504044), 0.1 mM NEAA, 2 mM l-glutamine, and 25 U/ml penicillin-streptomycin)] supplemented with 200 ng/ml of BMP4, 1000 U/ml of human LIF, 200 ng/ml of SCF, 100 ng/ml of EGF and 10 uM of Y-27632.

The floating aggregates were cultured for 8 days without replacing the medium.

**Generation of xrTestes aggregates**

Animal procedures were conducted in compliance with Institutional Animal Care and Use Committee of University of Pennsylvania. All mice were purchase from Charles River (Wilmington, MD).

xrTestes were generated by aggregating PGCLCs with fetal testicular somatic cells obtained from E12.5 mouse embryos as described previously (*2*). PGCLCs at d5 (humans) or d8 (rhesus macaque) of induction or after 2D expansion culture were used after FACS-sorting of AG$^+$ or EPCAM$^+$ITGA6$^+$ cells. To isolate fetal testicular somatic cells, E12.5 embryos were isolated from timed pregnant ICR or C57BL/6NCrSlc females mated with DBA/2CrSlc males and collected in chilled DMEM containing 10% FBS and 100 U/ml penicillin/streptomycin (Gibco). Fetal testes were identified by stripes, and the mesonephros were removed by tungsten needles.

Isolated testes were washed with PBS and then dissociated into single cells using 0.05% Trypsin-EDTA in PBS for 10 min at 37 °C followed by quenching with FBS. Cell suspensions were strained through a 70 μm nylon cell strainer and centrifuged. The remaining cell pellet was then resuspended with 100 μl MACS buffer (PBS containing 0.5% BSA and 2 mM EDTA) and incubated with anti-SSEA1 antibody MicroBeads (Miltenyi Biotec) for 20 min on ice before being washed with MACS buffer and centrifuged. The cell pellet was again resuspended in MACS buffer and then applied to an MS column (Miltenyi Biotec) according to the manufacturer's protocol. The flowthrough cells were centrifuged, resuspended with Cell Banker Type I, and cryopreserved in liquid nitrogen until use. All centrifugations were performed at 232 × g for 5 min and were followed by removal of the supernatant.

To generate floating aggregates, PGCLCs (5000 cells per xrTestis) and thawed fetal testicular somatic cells (60,000 cells per xrTestis) were mixed and plated in a Cell repellent V-bottom 96-well plate (Grener Bio-One, 651970) in Minimum Essential Medium alpha (α-MEM, Invitrogen) containing 10% KSR (Gibco), 55 μM 2-mercaptoethanol (Gibco), 100 U/ml penicillin/streptomycin (Gibco) and 10 μM Y-27632 for two days. For ALI culture, floating aggregates were transferred onto Transwell-COL membrane inserts (Corning, 3496) using a glass capillary. Membrane inserts were immersed in α-MEM supplemented as before, without Y-27632. xrTestes were cultured at 37 °C under an atmosphere of 5% $CO_2$ in air and one-half the volume of medium was changed every three days. For transplantation, floating aggregates were cultured for two days and transplanted.

**Transplantation of xrTestes aggregates under the kidney capsule of mice**

6 to 8 weeks female NOD-*Prkdc*$^{em26Cd52}$*Il2rg*$^{em26Cd22}$/NjuCrl (NCG) mice were used for transplantation. Mice were housed under a 12:12 h light: dark cycle at 22–24°C, with a humidity of 40–60%. Surgery was performed aseptically following the IACUC guidelines for rodent survival surgery. xrTestes aggregates were transplanted under the kidney capsules of bilaterally ovariectomized recipient female mice of NCG models. Mice were anesthetized by an intraperitoneal (IP) injection of Ketamine/Xylazine/Acepromazine

(100 mg/kg Ketamine; 10 mg/kg Xylazine; 2 mg/kg Acepromazine). Incision was made through the muscular body wall overlying the kidney, which was then exteriorized by applying pressure using the forefinger and thumb. After the initial hole was made in the delicate capsule, a pocket was created and enlarged using the fire-polished pipette. xrTestes aggregates were inserted into the pocket by lifting the edge of the kidney capsule and pushing the grafts into place with the fire-polished pipette. Once the kidney was returned into the peritoneal cavity, ovariectomy was performed by separating the musculature using curved tip scissors and carefully pulling the ovarian fat pad out of the same incision. Using the tweezers hemostatic, the region below the ovary was tightly clamped to be removed.

**Fluorescence-activated cell sorting (FACS)**

We isolated d5 PGCLCs used for xrTestis formation via FACSAria Fusion sorter. Floating aggregates at d5 containing PGCLCs derived from human iPSCs, were dissociated into single cells with 0.1% Trypsin/EDTA treatment for 15 min at 37 °C with periodic pipetting. After the reaction was quenched by adding an equal volume of FBS, cells were resuspended in FACS buffer (0.1% BSA in PBS) and strained through a 70 μm nylon cell strainer to remove cell clumps. $AG^+$ cells were FACS-sorted and collected in an Eppendorf tube containing α-MEM. For the analysis/sorting of PGCLCs with cell-surface markers, the cells dissociated with Trypsin-EDTA/PBS were stained with the fluorescence-conjugated antibodies (BV421-conjugated anti-CD49f and APC-conjugated anti-CD326) for 15 min at room temperature. After washing cells twice with the FACS buffer, the cell suspension was filtered by a cell strainer and analyzed or sorted by FACS.

For isolating germ cells from grafted xrTestes, xrTestes were carefully separated from the kidney by using forceps after confirmation of euthanasia. Isolated xrTestes were minced by scissors and dissociated by Multi Tissue Dissociation Kit 1 (Miltenyi Biotec) according to the manufacturer's protocol. xrTestes-derived germ cells were analyzed and sorted using a similar procedure, except 0.1% BSA fraction V (Gibco) in PBS was used to wash the cells prior to straining and cells in individual fractions (i.e., $AG^{+/-}VT^{+/-}PC^{+/-}$) cells were collected in CELLOTION (Amsbio). All FACS data were collected using FACSDiva Software v 8.0.2 (BD Biosciences).

**Immunofluorescence (IF) analysis**

Antibodies used for IF analysis is listed in Table S5. For IF analysis of xrTestes tissue, xrTestes were fixed with 4% paraformaldehyde (Sigma) in PBS for 2 h at RT, washed three time with PBS containing 0.2% Tween-20 (PBST), and then successively immersed in 10% and 30% sucrose (Fisher Scientific) in PBS overnight at 4 °C. The fixed tissues were embedded in OCT compound (Fisher Scientific), frozen, and sectioned to 10 μm thickness using a −20 °C cryostat (Leica, CM1800). Sections were placed on Superfrost Microscope glass slides (Thermo Fisher Scientific) that were then air-dried and stored at −80 °C until use.

Prior to staining, slides were treated with HistoVT one (Nacalai USA) for 20 min at 70 °C for the antigen retrieval. Subsequently, slides were washed three times with PBS and then incubated with blocking solution (5% normal goat serum in PBST) for 1 h. Slides were subsequently incubated with (1) primary antibodies in blocking solution for 2 h, followed by (2) secondary antibodies and 1 μg/ml DAPI in blocking solution for 50 min. Each incubation was performed at room temperature and followed by four washes with PBS (15 min each). Slides were mounted in Vectashield mounting medium (Vector Laboratories) for confocal laser scanning microscopy analysis (Leica, SP5-FLIM inverted). Confocal images were processed using Leica LasX (version 3.7.2).

For IF analyses of some of xrTestes, samples were fixed in 10% buffered formalin (Fisher Healthcare) with gentle rocking for 24 hr at room temperature. After dehydration, tissues were embedded in paraffin, serially sectioned at 4 μm thickness using a microtome (Thermo Scientific Microm™ HM325) and placed on Superfrost Microscope glass slides. Paraffin sections were then de-paraffinized using xylene. Antigens were retrieved by treating sections with HistoVT One for 35 min at 90 °C and then for 15 min at room temperature. The staining and incubation procedure for paraffin sections was similar to that for frozen sections, with the following modifications: the blocking solution was 5% normal donkey serum in PBST; the primary antibody incubation was performed overnight at 4 °C; and slides were washed with PBS six times after each incubation. Slides were mounted in Vectashield mounting medium for confocal microscopic analysis.

For IF of iPSCs, cells were cultured on MEFs plated 35 mm glass bottom dish. At d7 of culture, the cells were fixed in 4% paraformaldehyde in PBS for 15 min at room temperature, washed three times with PBS for 5 min each and incubated in 0.2% Triton-X100 (Fisher, BP151-100) in PBS for 10 min at room temperature. Then, the cells were incubated with primary antibodies in blocking solution for 1 h, followed by secondary antibodies and 1 μg/ml DAPI in blocking solution for 50 min. Images were captured and processed by an inverted microscope (Leica, DMi8).

For IF of rhesus macaque PGCLCs after expansion culture, cells were cultured on Glass Bottom Dish (Matsunami, D11130H) plated with mitomycin C-treated STO feeder cells. At d10 of culture, cells were fixed in 4% paraformaldehyde in PBS for 15 min at room temperature, washed three times with PBS (5 min each) and incubated in 0.2% Triton-X100 in PBS for 10 min at room temperature. Then, cells were incubated with primary antibodies in blocking solution for 1 h, followed by secondary antibodies and 1 μg/ml DAPI in blocking solution for 50 min. Both incubations were performed at room temperature and followed by four PBS washes. Images were captured and processed by confocal laser scanning microscopy.

**In situ hybridization**

ISH on formalin-fixed paraffin-embedded sections was performed using the ViewRNA ISH Tissue Assay Kit (Thermo Fisher) with gene-specific probe sets for human *PIWIL4* (VA1-3014459VT), human *MEIOB* (VA1-3016011), human *ACTB* (VA1-103510VT, positive control) and Bacillus subtilis *dapB* (VF1-11712VT, negative control). Experiments were performed according to the manufacturer's instructions.

**Quantitative PCR (qPCR) analysis**

For qPCR analysis, cDNA synthesis followed by total RNA extraction was performed as described previously (*8*). FACS-sorted rhesus macaque PGCLCs were collected in

CELLOTION. Rhesus macaque iPSCs were collected in the PBS without FACS sorting. Briefly, total RNAs were extracted from the cells using RNeasy Micro kits (QIAGEN, 74104) according to the manufacturer's instructions. The cDNA synthesis from 1 ng of total RNAs and the amplification of their 3′ends were performed as described previously (*8*). The quality of the amplified cDNAs was validated by examining the Ct values by qPCR with the primers listed in Table S6. Quantitative PCR was performed using Power SYBR Green PCR Master Mix (Thermo Fisher, 4367659) with a StepOnePlus real-time qPCR system (Applied Biosystem) according to the manufacturer's instructions. For each gene examined, the ΔCt from the average Ct values of the two independent housekeeping genes *GAPDH* and *PPIA* (set as 0) were calculated and plotted for two independent experiments. Mean ± SD of three technical replicates are shown.

**10x Genomics single-cell RNA-seq library preparation**

xrTestis-derived cells were sorted via FACS for expression of AG or VT as described previously and collected in CELLOTION(*2*). Cell clumps were removed by straining cell suspensions through a 70 μm nylon mesh and then the single-cell suspension was centrifuged at 300 × g for 5 min. Cells were resuspended in 0.1% BSA in PBS and counted. All samples had greater than 80% viability as determined by trypan blue staining. Encapsulation was performed using a 10x Genomics Chromium Controller according to the manufacturer's protocol. The cells were loaded into Chromium microfluidic chips with the Chromium Single Cell 3' Reagent Kit (v3.1 chemistry) and then used to generate single-cell Gel Bead emulsions (GEMs) using the Chromium Controller (10x Genomics) according to the manufacturer's protocol. Reverse transcription step was performed using a C1000 Touch Thermal Cycler with 96-Deep Well Reaction Module (Bio-Rad). All subsequent cDNA amplification and library construction steps were performed according to the manufacturer's protocol. Libraries were sequenced using a 150-cycles paired-end sequencing protocol on a HiSeq 4000 or NovaSeq 6000 instrument.

**Read Mapping of scRNA-seq data and downstream analysis**

Sequence data was demultiplexed and fastqs produced using Cell Ranger (v.7.1.0). For human samples, reads were mapped to the GRCh38 human reference and independently mapped to the GRCm38 mouse reference. Similarly, macaque samples were aligned to the Mmul_10 reference, created using the Mmul_10 genome assembly and gtf file using Cell Ranger's mkref function, as well as to the mouse reference. This produced two separate alignments for each cell, one for the human/macaque reference (as appropriate for human or macaque samples) and another for the mouse reference. Both sets of UMI counts were read into R (v.4.2.2) and secondary analysis was performed using Seurat (v.4.4.0) (*9*). UMI count tables were first loaded into R by using the Read10x function, and Seurat objects were built from each sample, one per alignment. The sum of UMI counts for each alignment was totaled for each cell for both human/macaque and mouse counts. The total UMIs for human/macaque alignment were then compared with the mouse alignment. Any cell with greater than twofold assignment of UMIs to human/macaque than mouse was designated as human or macaque as appropriate for the sample. Similarly, any cell with twofold higher mouse counts than primate would be designated mouse. Any cell that failed to meet either threshold was labeled "undetermined" and excluded from the analysis. Human (iPSCs and *in vivo* 12-year-old testis) samples and mouse (two independent adult testes) samples were used as controls. No human or mouse control cells were assigned to be the wrong species (Fig. S4A). In this fashion, all cells with "mouse" or "undetermined" identities were excluded, and only human cells aligned to the human reference or macaque cells aligned to the macaque reference were kept for downstream analysis. For each sample a minimum number of genes = 100 and maximum genes = 3000 were set as well as a maximum mitochondrial content of 20% and cells outside those bounds were excluded. A minimum UMI was set on a per-sample basis to exclude low-quality cells by assigning a minimum cutoff based on an inspection of a ranked plot of UMI per sample. For the sample NCG8, a cutoff of $10^{3.5}$ was set, for NCG5, NCG19 and NCG_c41 a cutoff of $10^{3.7}$ was used, while for NCG4, NCG9 and NCG_182_2 a cutoff of $10^{3.8}$ was applied, and for all other samples a minimum of $10^{4}$ UMI/cell was set. Samples were integrated using Seurat's IntegrateData function using 3000 highly-variable genes and the first 30 dimensions. Cell cycle state was assigned using Seurat's CellCycleScoring function and samples were split by phase and re-integrated to regress out cell cycle effects. Doublets were identified using DoubletFinder (*10*), with an expected doublet rate of 4%. This identified three clusters which showed a high rate of doublets and with an intermediate identity when looking at key marker genes, so these clusters were removed, and the dataset was re-clustered using 30 principle components. Following these filtering steps, 10,298 cells were brought forward for further analysis, grouped via Seurat into 18 clusters. RNA velocity was determined using velocyto (*11*) with the parameters deltaT = 10, kCells = 25, fit.quantile = 0.01. Pseudotime was generated using monocle3 (ref). As PGCLCs showed a clear disjunction from the unbroken progression of all other germ cells due to a lack of intermediate cell types captured, they were excluded before running pseudotime, which was generated using monocle's learn_graph function with ncenter=50, minimal_branch_len=1. Using previously identified marker genes for PGCs, prospermatogonia and postnatal germ cell types, clusters were assigned to one of ten cell types, ranging from early PGCs to preleptotene spermatocytes. A random selection of *MEIOB* transcripts were selected by searching fastq files for a 3' *MEIOB* sequence and aligned to the human *MEIOB* and mouse *Meiob* genes, and the cells containing these transcripts were identified via cell barcode. These showed 100% match to human *MEIOB* and only 35% identity with mouse *Meiob* via BLAST for the 3' region of the gene. When the cells that contain these human aligned *MEIOB* molecules were mapped back onto the UMAP, they corresponded to the region of *MEIOB* expression.

DEGs between groups were determined using Wilcoxon Rank Sum test via Seurat's FindMarkers function. For pseudobulk analysis, mean normalized counts were used by cell type. Gene ontology enrichment was analyzed via DAVID (v2023q4) (*12*). Ingenuity Pathway Analysis (QIAGEN) was used for all pathway analyses.

To examine the heterogeneity of the spermatogonial compartment, T1 prospg, spg and Diff. spg.E were subset, reclustered, and pseudotime run as described above. Four clusters were identified that corresponded to T1 prospg, S0 Spg, S1 Spg and Diff. spg.E.

To compare xrTestis with normal *in vivo* spermatogenesis we used an atlas of human germ cells we have derived previously (*13*). Briefly, testis cells derived from between gestational week (GW) 7 and GW 17 fetuses, along with cells from individuals from shortly after birth to 12 years old were analyzed via scRNA-seq. *In vivo* samples were first integrated together in the same manner as described for xrTestes. Somatic cells, identified by key gene expression, were eliminated and the remainder reclustered and cell cycle phase state assigned via Seurat as described previously. *In vivo* germ cells were then split by cell cycle phase and integrated with xrTestis data, also split by phase. The integrated object was clustered by Monocle 3 and a UMAP generated by Seurat and cell types were assigned to these new clusters anew, replacing prior designations. Similarities and differences between *in vivo* and xrTestis clusters were therefore based on shared clusters assigned together. Differentially expressed markers were defined using FindMarkers and pseudobulk DEGs as described previously.

**scRNA-seq analysis of rhesus macaque cells**

Analysis of macaque xrTestis-derived germ cells was performed in the same manner as human germ cells. For comparison of macaque and human datasets, data transfer was employed using macaque dataset as a query against the human datasets, using a subset of homologous genes with identical genes. Seurat's MapQuery and TransferData functions were used to project macaque cells onto human dimensional reduction/UMAP.

**Whole genome bisulfite sequencing**

PGCLC aggregates were digested with 400 µl 0.25% trypsin for 15 min. Then, 100 µl

FBS was used to stop digestion, and the lysates were pipetted well to obtain a single-cell suspension. The fractions of $AG^{+}VT^{-}$, $AG^{+}VT^{+}$ and $AG^{-}VT^{+}$ cells were FACS-sorted for methylome analyses. Cells were collected and lysed in 50 mM Tris (pH 8.0), 10 mM EDTA, 0.5% SDS, and 100 μg/ml proteinase K. Then, crude DNA was used to build a sequencing library according to the protocol of the Pico Methyl-Seq Library Prep Kit (Zymo, #D5455). The libraries were sequenced on the Illumina 2200 platform. Raw fastq files were demultiplexed with bcl2fastq2 (v.2.20). Barcode and index trimming was performed with Trim Galore (v.0.6.5) as follows: trim_galore --quality 30 --phred33 --illumina --stringency 1 --cores 4 -e 0.1 --fastqc --clip_R1 10 --three_prime_clip_r1 10 --length 20. The trimmed fastq files were then mapped to calJac4 from USCS with Bismark (v.0.22.3) as follows: bismark --parallel 4 --genome_folder $REF --non_directional --score_min L,0,–0.6. CpG methylation was extracted and analyzed with methylKit (v.1.22.0). Covered CpG loci were included in the analysis. The genome was tiled in 2 kb windows, and DNA methylation levels were summarized with methyKit (v.1.22.0). Data were visualized with R (v.4.1.0).

**Supplementary Figure Legends**

**Supplementary Figure 1. xrTestes display markers of mouse somatic cells and human germ cells.**

(A) Fluorescence-activated cell sorting (FACS) plot of d5 PGCLCs derived from 9A13 human iPSCs. The percentage of AG (+) cells in the indicated gate is shown.

(B) (Left) Bright field images of d2 floating xrTestis aggregates before transplantation. Scale bar, 200 µm. (Right) FACS plot of d2 floating aggregates to assess the number of AG (+) PGCLC-derived cells per aggregate.

(C) BF images of xrTestis at 1 month post-transplantation grafted under the murine kidney capsule. Scale bars: 500 µm.

(D) IF images of reconstituted (r)Testis or xrTestes at 1 month post-transplantation. Markers for PGCLC-derived cells (AG, TFAP2C, SOX17), Sertoli cell marker (SOX9), and a basement membrane marker (LAMININ), with their merges with DAPI staining (white). Scale bars, 20 µm. rTestes are made with mouse testicular somatic cells without human components.

(E) IF images of 1month xrTestes for GC markers (AG, NANOG, or POU5F1), a Sertoli cell marker (SOX9), and somatic cell markers (HSD3B [Leydig cell marker], NR2F2 [stromal cell marker], ACTA2 [peritubular myoid cell/perivascular smooth muscle cell marker], CD31 [vascular endothelial marker]) with their merges with DAPI staining (white). Scale bars, 20 µm.

**Supplementary Figure 2. Expansion culture of human PGCLCs**

(A) FACS-sorted AG (+) human PGCLCs were cultured in a medium containing forskolin, SCF and bFGF (see methods) and passaged every 7-10 days, sorting for AG (+) cells.

(B) BF images of c37, c39, c41 (at passage 5) and c90 (at passage 12) colonies of human PGCLCs. Scale bars, 100 μm.

(C) FACS plots showing AG expression of PGCLCs at d5 and at c10 to c90 of expansion culture.

(D, E) The percentages of AG (+) cells (D) and growth curves of AG (+) cells as indicated by $\log_{10}$ fold increases (E) during the expansion culture until c41 and c90. Circle or square dot represents each replicate.

(F, G) (left) BF images (inset) and FACS plots of d2 floating xrTestis aggregates using c41 (F) and c90 PGCLCs (G). Scale bar, 200 μm. The number of AG (+) PGCLC-derived cells per aggregate are shown. (middle) Bright field images of d120 xrTestis using c41 PGCLCs (F) and d180 xrTestis using c90 PGCLCs (G), scale bar, 500 μm. (right) FACS plots of xrTestes.

**Supplementary Figure 3. Derivation of human iPSCs carrying *TFAP2C-EGFP;DDX4-tdTomato* reporter alleles**

(A) FACS analysis of d5 PGCLC derived from 679SeV4 iPSCs stained with EPCAM and ITGA6. The percentage of cells in the indicated gate is shown.

(B) xrTestis derived from 679SeV4 iPSCs at d120 of transplantation, scale bar, 500 μm.

(C) IF images of d120 679SeV4 xrTestis highlighting PGCLC-derived cells (green:

MAGEC2, red: DDX4, cyan: hMito) with their merges with DAPI staining (white). Scale bars 20 µm.

(D) Schemes of the human *TFAP2C* and *DDX4* loci and the targeting constructs for generating *TFAP2C-p2A-EGFP*; *DDX4-p2A-tdTomato* alleles. In both cases, selection cassettes were excised by Cre recombination. Black boxes indicate exons. Arrows show the positions of the genotyping primers.

(E) Genotyping PCR for 18-2 AGVT iPSCs confirming heterozygous and homozygous knock-in of reporter genes for the *TFAP2C* and *DDX4* loci, respectively. Negative control: 679Sev4 iPSCs; positive control: 9A13 iPSCs. Rec, recombined allele; non, non-recombined allele.

(F) Phase contrast image of 18-2 iPSCs, scale bar, 200 µm.

(G) FACS plot of d5 PGCLCs derived from 18-2 iPSCs.

(H) FACS plot of floating aggregates on day 2 (mixed aggregates of $AG^+$ FACS-sorted PGCLCs and mouse testicular somatic cells, immediately before transplantation). Inset: brightfield view of an aggregate, scale bar: 200 µm.

(I) BF images (left) and FACS plot (right) of xrTestes derived from 18-2 iPSCs after 120 days of transplantation. Scale bars, 500 µm. The percentage of cells in the indicated fractions are shown.

**Supplementary Figure 4. Gene expression changes throughout xrTestis germ cell differentiation**

**(A)** Proportion of cells with identity matched to human, mouse and undetermined.

Control mouse testis cells, human iPSCs and human testis cells were used to confirm 0% misattribution of cells to the wrong species by the algorithm.

(B) Unbiased clustering of xrTestis germ cells projected onto UMAP. Cell type assignments to clusters are indicated by dotted lines.

(C) RNA velocity projected onto clusters.

(D) Pseudotime generated by first excluding PGCLCs and using PGCs Late1 as the origin.

(E) Gene expression for xrTestis cells for a selection of germ cell markers. Cells are arranged along pseudotime and colored per cell types in Fig. 2A. Mean expression is shown as a black line.

(F) Violin plots of gene expression for select markers by cell type.

(G) Ridgeplot showing frequency of germ cells over pseudotime within scRNAseq datasets, broken up by sample and organized by transplant duration. Original iPSC line is shown on the right.

(H) UMAP plots of scRNA-seq samples, colored by cell type assignment and arranged by time transplanted. iPSC line of origin is shown below. Samples maintained in 2D expansion culture before transplantation are shown on the right of the panel.

**Supplementary Figure 5. Gene expression changes throughout xrTestis germ cell differentiation.**

Comparison of genes expressed in pairwise progression of cell types. For each scatter plot, $log_2$ fold change > 0.25 and adjusted $p$ value < 0.05 were used to determine DEGs, colored by cell type matching the UMAP in Fig. 2. Select markers of interest have been labelled. Two representative GO terms have been selected for each DEG set and key genes involved in that process are shown.

**Supplementary Figure 6. Methylation status of xrTestis-derived germ cells**

5-methylcytosine (5mC) levels analyzed by whole genome bisulfite sequencing (WGBS) for PGCLCs and for three fractions of xrTestis germ cells selected using the gating strategy shown in Fig. 1F.

**Supplementary Figure 7. NANOS3 and DND1 are required for production of iPSC-derived germ cells.**

**(A)** Scatter plots comparing the average gene expression between PGCs.E and PGCs.L1. DEGs show greater than 4-fold expression change and $p_{adj} < 0.05$ are colored. Key DEGs including *NANOS3* and *DND1* are highlighted.

(B) Co-IF/ISH images of a human embryo section (4 wpf) for TFAP2C (green) and NANOS3 (red) merged with DAPI staining (white). Migrating TFAP2C$^{+}$ PGCs within the dorsal wall of the mesentery expressing *NANOS3* RNA are shown. Scale bar, 20 µm.

(C) Scheme of air-liquid interface (ALI) culture of xrTestes using PGCLCs derived from wild-type or mutant iPSC lines (NANOS3-KO and DND1-KO).

(D) Violin plot showing the expression of *DND1* and *NANOS3* from scRNA-seq data obtained from the indicated cell types.

(E) FACS plots of d5 PGCLC-containing aggregates induced from wild-type (WT) or

NANOS3 mutant (NANOS3 KO) iPSCs. Percentages of AG$^+$ PGCLCs are shown.

(F) Phase-contrast images of d14 xrTestes. Scale bar, 200 µm.

(G) IF images of d14 xrTestis (WT or NANOS3 KO) sections for TFAP2C-EGFP (green), TFAP2C (red) and SOX9 (cyan) merged with DAPI staining (white). Scale bar, 20 µm.

(H) FACS plots of xrTestes derived from WT and *NANOS3* KO lines after three weeks of ALI culture. The total number of AG$^+$ germ cells per xrTestis are shown.

(I) The total number of AG$^+$ germ cells per xrTestis at indicated time points as assessed by FACS (n =3–4). Means ± standard deviation.

(J) The total number of AG$^+$ germ cells per xrTestis at indicated time points. Germ cells were induced from indicated iPSC lines. WT, wild-type parental iPSCs; KO+OE, *NANOS3* mutant iPSCs bearing dox-inducible NANOS3 over-expression cassette; DOX, doxycycline. Means ± standard deviation

showing rescue of the *NANOS3*-KO by doxycycline-inducible overexpression vector (dox). Error bars denote standard deviation.

(K) IF of xrTestis sections (WT and DND1 KO, cultured by ALI for 14 days) for GFP (TFAP2C-EGFP, green), TFAP2C (red) and SOX9 (cyan). Scale bar, 50 µm.

(L) The total number of AG$^+$ germ cells per xrTestes as in (K). Means ± standard deviation. $p<0.05$.

(M) Bright field images of xrTestis xenografts under the kidney capsule (3 weeks after transplantation). Scale bar, 1 mm.

(N) FACS plot of xrTestes as in (K). The total number of AG$^+$ germ cells per xrTestis are shown.

(O) The total number of $AG^+$ germ cells per xrTestis as in (N). Means ± standard deviation. Statistical significance assessed by Welch's *t*-test, $p<0.05$.

(P) Percentage of cells expressing TFAP2C-EGFP among all human mitochondrial antigen (hMito) expressing cells in xrTestes derived from WT and *NANOS3* KO iPSCs as assessed by IF.

(Q) IF of xrTestis sections for hMito (red), TFAP2C (green), NANOG (cyan) and DAPI staining (white). Scale bar, 20 μm.

(R) Scatter plot comparing the average expression values (from scRNA-seq) between WT and *NANOS3* KO $AG^+$ cells in d9 xrTestes. DEGs ($log_2FC > 0.25$ and $p_{adj} < 0.05$) are highlighted in colors. Representative genes and their GO enrichments upregulated in mutant cells are shown at right.

(S) Scatter plot comparing the average expression between WT and *DND1* KO $AG^+$ cells in d14 xrTestes.

**Supplementary Figure 8. IF of xrTestes for key stage-specific germ cell markers.**

IF of xrTestes at 6 months after transplantation for indicated proteins. Merged images with DAPI staining (white) are shown at right. Scale bars: 20 μm.

**Supplementary Figure 9. Comparison of *in vivo* and xrTestis germ cells**

**(A)** Expression of key markers throughout spermatogenesis projected on the UMAP plot as in Fig. 4B.

(B) Pearson correlation of gene expression between *in vivo* and xrTestis cell types.

(C) Ingenuity pathway analysis of DEGs of indicated cell types (in vivo and xrTestis). *Z*-score of activation/repression of the pathway is shown by color, significance of overlap between cell-type-specific genes and pathway genes is indicated by size of the dot (-$\log_{10}$ *p*-value).

**Supplementary Figure 10: Comparison of *in vivo* versus xrTestis cell types**

Scatter plots comparing the average gene expression of the indicated cell types between in vivo and those derived from xrTestes. DEGs ($\log_2$ fold change > 0.25 and adjusted *p*-value < 0.05) are highlighted with colors (red, upregulated in vivo; blue, upregulated in xrTestis). Representative genes and their GO enrichments for DEGs are shown beneath each plot.

**Supplementary Figure 11. Identification of two distinct subsets of spermatogonia in both in vivo and xrTestes.**

(A) UMAP plot showing two distinct subsets of spg (S0 and S1 spg) after reclustering of “T1 prospg”, “Spg” and “Diff. spg.E” from Fig. 4A. Pseudotime is shown overlaid with a direction as indicated by the arrow.

(B) Human *in vivo* cells from UMAP plot in (A). Inset pie chart showing proportion of S1 and S0 spg.

(C) Human xrTestis cells from UMAP plot in (A). Inset pie chart showing proportion of S1 and S0 spg.

(D) Expression of representative marker genes projected on the UMAP plot as in (B, in vivo) and (C, xrTestis).

(E) Pearson correlation matrix showing similarity between xrTestis and *in vivo* cell types.

(F) Scatter plots comparing the average gene expression between S0 and S1 spg for *in vivo* (left) and xrTestis (right). DEGs (0.3 < log2 fold change, adjusted *p*-value < 0.05) are highlighted in colors and key markers are indicated. Representative genes and their GO enrichments are shown beneath each plot.

(G) Projection of DEGs of in vivo cell types as defined in (F, left, highlighted in colors) on scatter plot comparing S0 and S1 spg from xrTestis as in (F, right).

(H) Heatmaps of indicated datasets showing the average expression of genes associated with the GO:0005743~mitochondrial inner membrane term significant in both datasets.

**Supplementary Figure 12. Establishment of xrTestis in rhesus macaque**

(A) Schematic illustration of the rhesus macaque TFAP2C and DDX4 loci and targeting construct for generating *AP2γ/TFAP2C-p2A-EGFP* (AG); *VASA/DDX4-p2A-tdTomato* (VT) alleles. Black boxes, coding sequences; grey boxes, 3' untranslated regions (3' UTR); red arrows, positions of PCR primers for genotyping.

(B) Genotyping PCR to screen for clones bearing AG (top) and VT (middle) alleles without random integration (bottom). The positions of the DNA fragments corresponding to AG with the pgk-Puro cassette excised and to VT with the pgk-Neo cassette excised, and the wild-type sequences (WT) are indicated. Flanking lanes are size markers.

(C) DNA sequences flanking the guide RNAs of the wild-type alleles for the *TFAP2C* and *DDX4* loci.

(D) Representative results of G-band karyotype analysis of 1C2AGVT19 iPSCs (i.e., clone 19 in B) bearing AG and VT alleles. Cells displayed a normal male karyotype (42, XY).

(E) Phase contrast images of 1C2AGVT19 iPSCs. Scale bar, 200 μm.

(F) IF images of 1C2AGVT19 iPSCs, stained as indicated. Scale bar, 50 μm.

(G) BF (top) and fluorescence images (bottom, green) of floating aggregates during PGCLC induction at indicated days. Scale bars, 250 μm.

(H) FACS plot of floating aggregates at d8 of PGCLC induction derived from 1C2AGVT19 iPSCs. The percentage of $AG^+$ cells is shown.

(I) qPCR quantification of the indicated markers and housekeeping genes of $AG^+$ FACS-sorted d8 PGCLCs. ΔCt values were calculated by subtraction of the raw Ct values of each gene (mean value of two biological replicates) from the averaged Ct values of the housekeeping genes *GAPDH* and *PPIA*. Mean ± SD of two biological replicates is shown.

**Supplementary Figure 13. Expansion culture of rhesus macaque PGCLCs**

(A) Scheme for expansion culture of PGCLCs induced from rhesus macaque 1C2AGVT19 iPSCs.

(B) Phase contrast images of expansion culture day (c)2, c4, c6, c8 and c10 of PGCLC colonies. The white dashed lines outline PGCLC colonies. Scale bars, 250 μm.

(C) FACS plot of c10 PGCLCs.

(D) IF images of c10 PGCLCs for DAPI (white), GFP (green), TFAP2C (red) and NANOG (cyan), and the merged image. Scale bar, 25 μm.

(E) FACS analyses of c10, c23, c34, c45, c55, c65 and c75 PGCLCs. The percentages of TFAP2C-EGFP$^+$ cells are shown.

(F) Growth curve of TFAP2C$^+$ cells during rhesus macaque PGCLC expansion culture until c45. A total of 10,000 TFAP2C$^+$ d8 PGCLCs were used as a starting cell population. Means ± standard deviation of three biological replicates.

(G) qPCR quantification of the indicated markers during PGCLC expansion culture. Mean values are connected by a line.

(H) Teratoma formation in xrTestes xenograft 1 month after transplantation. d8 PGCLCs without expansion culture was used to generate xrTestes. Scale bar, 1 mm.

(I) xrTestes recovered from mice one month after transplantation. These xrTestis were made with c10, c23, c34 or c45 PGCLCs with expansion culture. Yellow arrow indicates teratoma. Scale bar: 1 mm.

(J) % occurrence of teratoma 1 month after xrTestis transplantation by culture time of PGCLCs through expansion culture.

**Supplementary Tables**

**Table S1. Differentially expressed genes (top 3000 genes) across cell types derived from human iPSCs.**

**Table S2. Differentially expressed genes by pairwise comparison between human iPSC-derived germ cell types**

**Table S3. Differentially expressed genes between in vivo germ cell types and corresponding iPSC-derived germ cell types**

**Table S4. Differentially expressed genes (top 3000 genes) across cell types derived from rhesus macaque iPSCs.**

**Table S5. Antibodies used in this study**

**Table S6. Primers used in this study**

### Figure S1

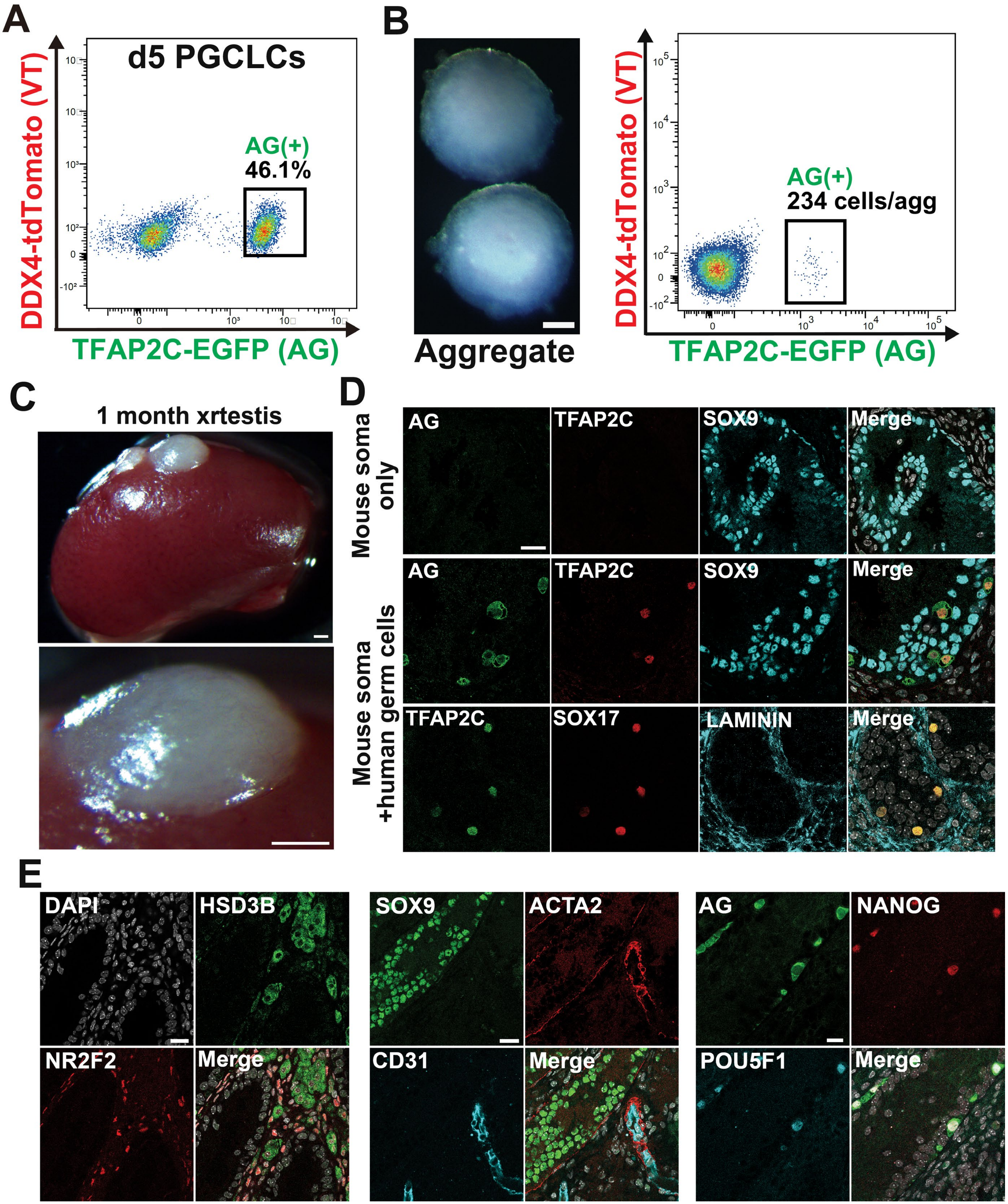
A
d5 PGCLCs
AG(+)
46.1%
DDX4-tdTomato (VT)
TFAP2C-EGFP (AG)
B
Aggregate
AG(+)
234 cells/agg
DDX4-tdTomato (VT)
TFAP2C-EGFP (AG)
C
1 month xrtestis
D
Mouse soma only
Mouse soma +human germ cells
AG
TFAP2C
SOX9
Merge
SOX17
LAMININ
E
DAPI
HSD3B
NR2F2
Merge
SOX9
ACTA2
CD31
AG
NANOG
POU5F1

### Figure S2

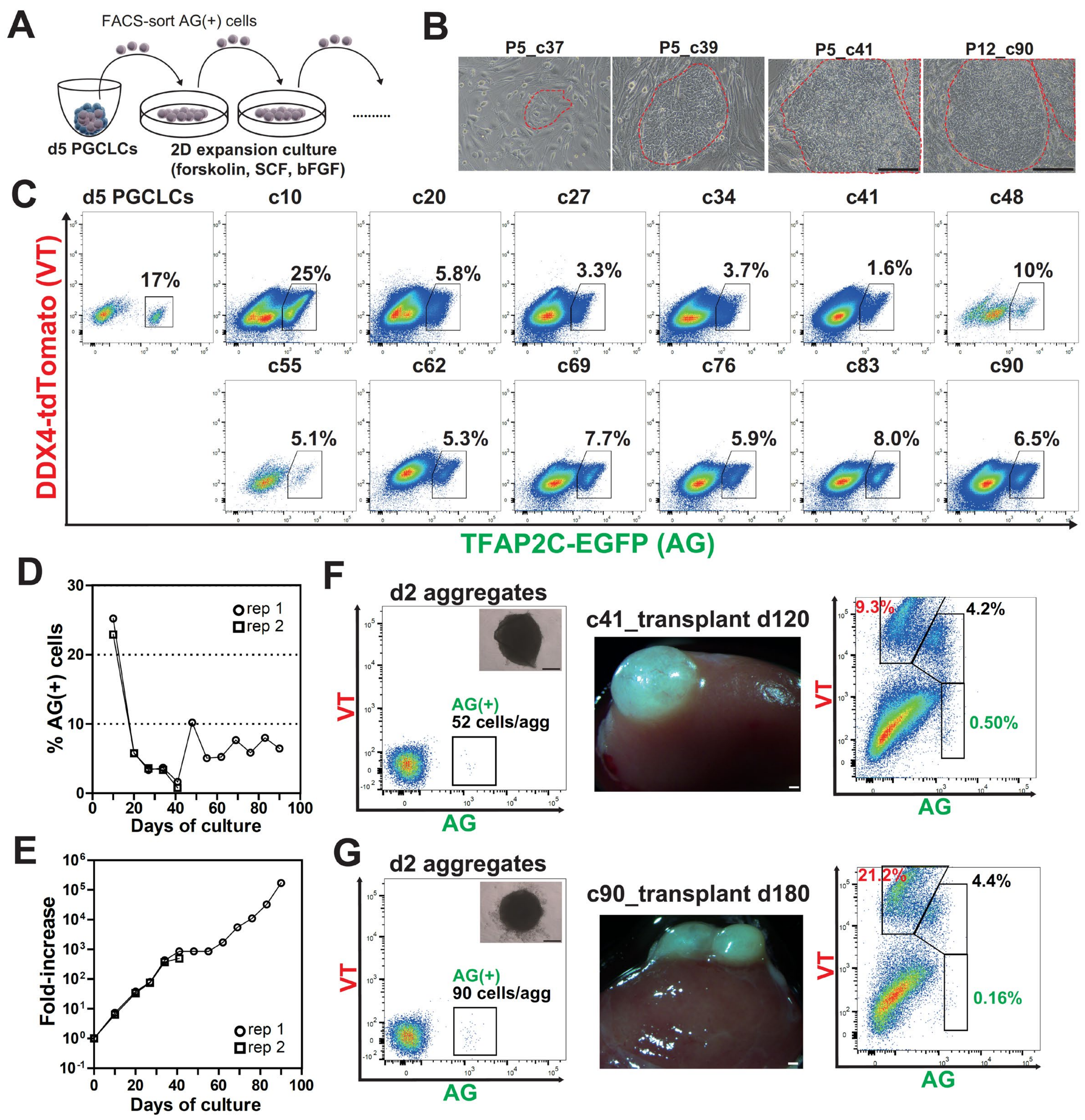
A
FACS-sort AG(+) cells
d5 PGCLCs
2D expansion culture
(forskolin, SCF, bFGF)
B
P5_c37
P5_c39
P5_c41
P12_c90
C
d5 PGCLCs
c10
c20
c27
c34
c41
c48
17%
25%
5.8%
3.3%
3.7%
1.6%
10%
c55
c62
c69
c76
c83
c90
5.1%
5.3%
7.7%
5.9%
8.0%
6.5%
DDX4-tdTomato (VT)
TFAP2C-EGFP (AG)
D
% AG(+) cells
rep 1
rep 2
Days of culture
E
Fold-increase
rep 1
rep 2
Days of culture
F
d2 aggregates
VT
AG(+)
52 cells/agg
AG
c41_transplant d120
9.3%
4.2%
0.50%
VT
AG
G
d2 aggregates
VT
AG(+)
90 cells/agg
AG
c90_transplant d180
21.2%
4.4%
0.16%
VT
AG

### Figure S3

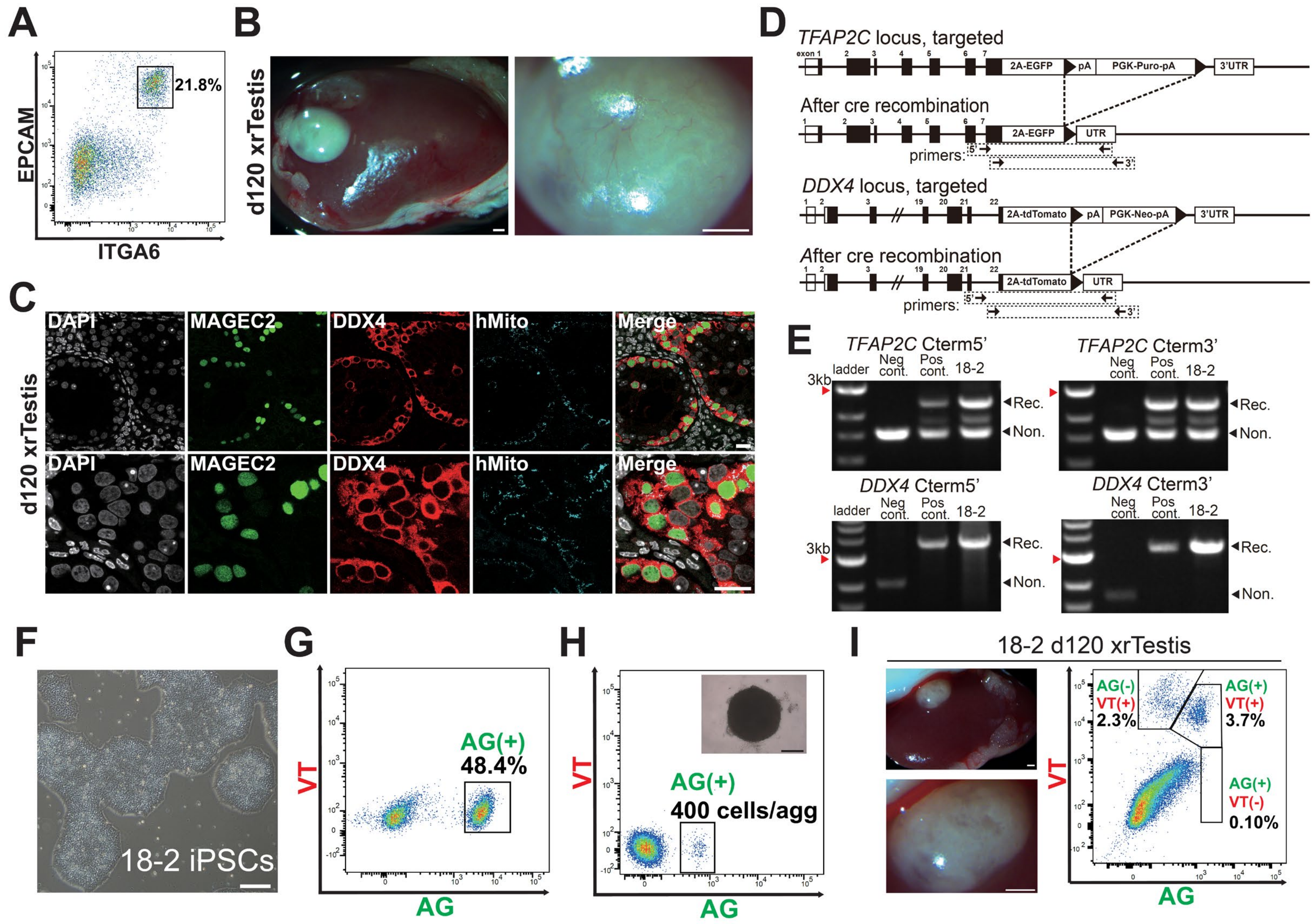
A
EPCAM
ITGA6
21.8%
B
d120 xrTestis
C
d120 xrTestis
DAPI
MAGEC2
DDX4
hMito
Merge
D
TFAP2C locus, targeted
exon 1
2A-EGFP
pA
PGK-Puro-pA
3'UTR
After cre recombination
UTR
primers:
DDX4 locus, targeted
2A-tdTomato
PGK-Neo-pA
E
TFAP2C Cterm5'
TFAP2C Cterm3'
DDX4 Cterm5'
DDX4 Cterm3'
ladder
Neg cont.
Pos cont.
18-2
3kb
Rec.
Non.
F
18-2 iPSCs
G
VT
AG
AG(+)
48.4%
H
AG(+)
400 cells/agg
I
18-2 d120 xrTestis
AG(-)
VT(+)
2.3%
AG(+)
VT(+)
3.7%
AG(+)
VT(-)
0.10%

### Figure S4

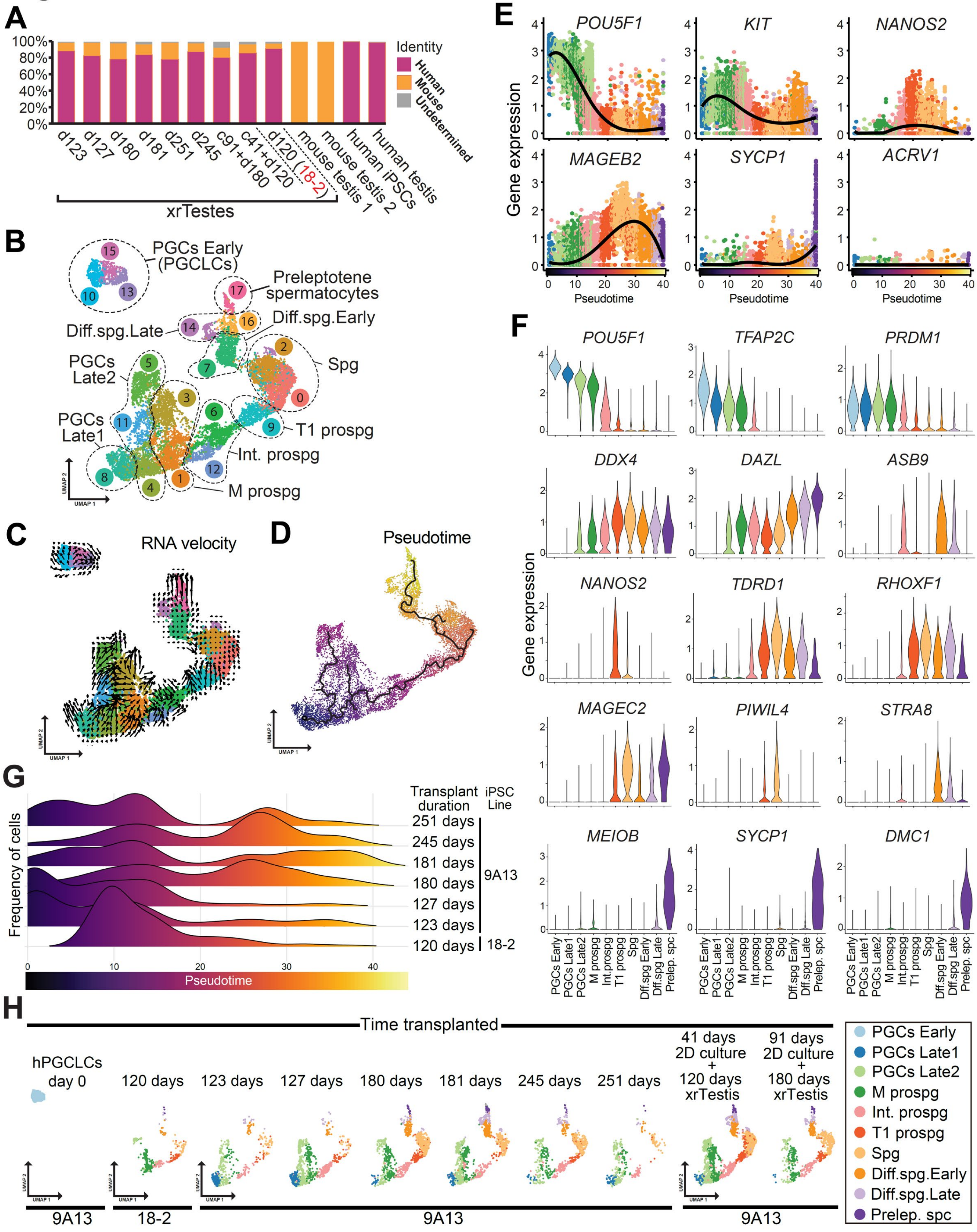
A
Identity
Human
Mouse
Undetermined
d123
d127
d180
d181
d251
d245
c91+d180
c41+d120
d120 (18-2)
mouse testis 1
mouse testis 2
human iPSCs
human testis
xrTestes
B
PGCs Early (PGCLCs)
Preleptotene spermatocytes
Diff.spg.Late
Diff.spg.Early
Spg
PGCs Late2
PGCs Late1
T1 prospg
Int. prospg
M prospg
C
RNA velocity
D
Pseudotime
E
Gene expression
POU5F1
KIT
NANOS2
MAGEB2
SYCP1
ACRV1
Pseudotime
F
POU5F1
TFAP2C
PRDM1
DDX4
DAZL
ASB9
NANOS2
TDRD1
RHOXF1
MAGEC2
PIWIL4
STRA8
MEIOB
SYCP1
DMC1
Gene expression
PGCs Early
PGCs Late1
PGCs Late2
M prospg
Int.prospg
T1 prospg
Spg
Diff.spg Early
Diff.spg Late
Prelep. spc
G
Frequency of cells
Transplant duration
iPSC Line
251 days
245 days
181 days
180 days
127 days
123 days
120 days
9A13
18-2
Pseudotime
H
Time transplanted
hPGCLCs day 0
120 days
123 days
127 days
180 days
181 days
245 days
251 days
41 days 2D culture + 120 days xrTestis
91 days 2D culture + 180 days xrTestis
9A13
18-2
9A13
9A13
PGCs Early
PGCs Late1
PGCs Late2
M prospg
Int. prospg
T1 prospg
Spg
Diff.spg.Early
Diff.spg.Late
Prelep. spc

### Figure S5

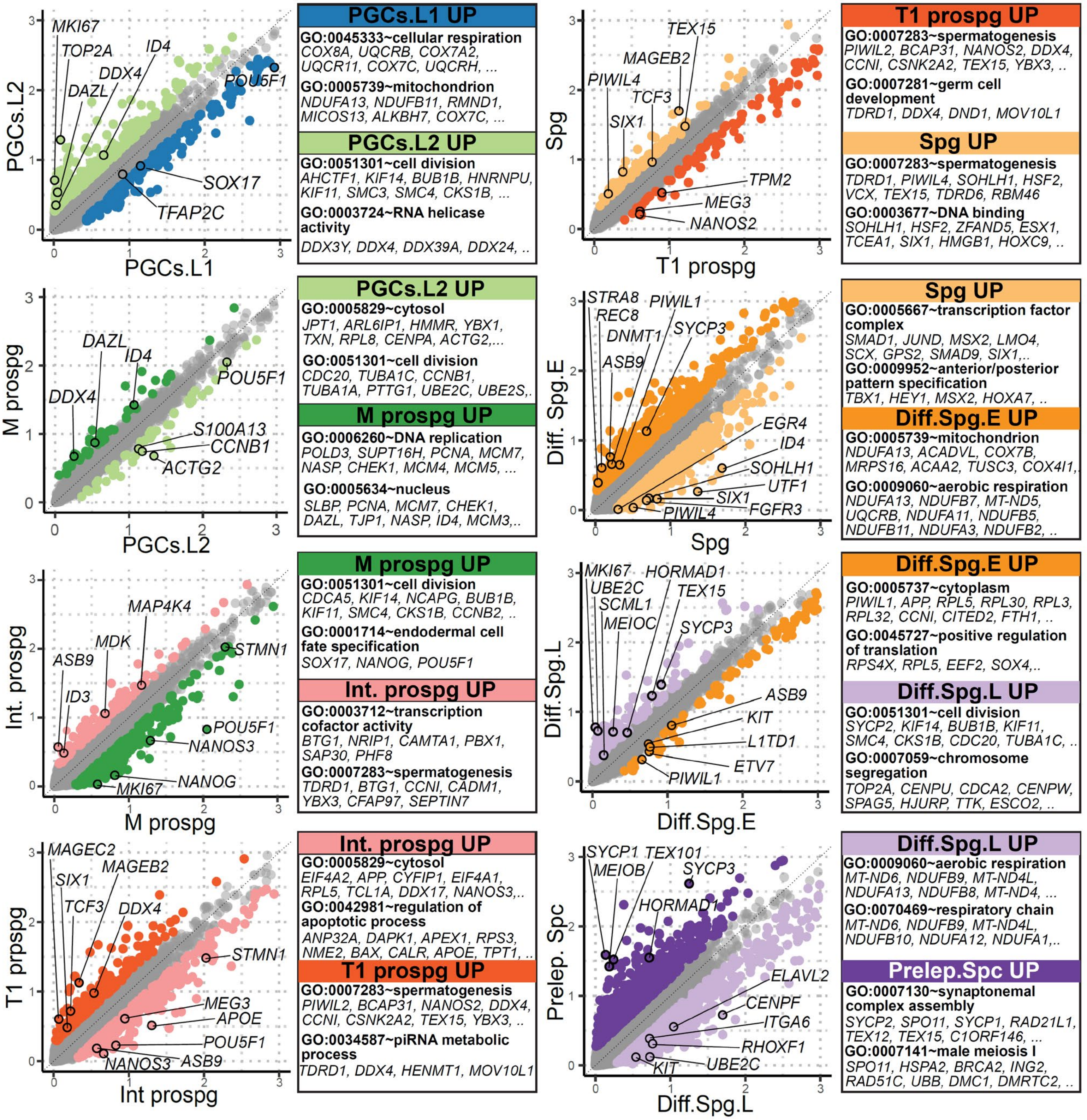

PGCs.L1
PGCs.L2
MKI67
TOP2A
ID4
DDX4
DAZL
POU5F1
SOX17
TFAP2C
PGCs.L1 UP
GO:0045333~cellular respiration
COX8A, UQCRB, COX7A2, UQCR11, COX7C, UQCRH, ...
GO:0005739~mitochondrion
NDUFA13, NDUFB11, RMND1, MICOS13, ALKBH7, COX7C, ...
PGCs.L2 UP
GO:0051301~cell division
AHCTF1, KIF14, BUB1B, HNRNPU, KIF11, SMC3, SMC4, CKS1B, ...
GO:0003724~RNA helicase activity
DDX3Y, DDX4, DDX39A, DDX24, ..
T1 prospg
Spg
TEX15
MAGEB2
PIWIL4
TCF3
SIX1
TPM2
MEG3
NANOS2
T1 prospg UP
GO:0007283~spermatogenesis
PIWIL2, BCAP31, NANOS2, DDX4, CCNI, CSNK2A2, TEX15, YBX3, ..
GO:0007281~germ cell development
TDRD1, DDX4, DND1, MOV10L1
Spg UP
GO:0007283~spermatogenesis
TDRD1, PIWIL4, SOHLH1, HSF2, VCX, TEX15, TDRD6, RBM46
GO:0003677~DNA binding
SOHLH1, HSF2, ZFAND5, ESX1, TCEA1, SIX1, HMGB1, HOXC9, ..
PGCs.L2
M prospg
DAZL
ID4
DDX4
POU5F1
S100A13
CCNB1
ACTG2
PGCs.L2 UP
GO:0005829~cytosol
JPT1, ARL6IP1, HMMR, YBX1, TXN, RPL8, CENPA, ACTG2,...
GO:0051301~cell division
CDC20, TUBA1C, CCNB1, TUBA1A, PTTG1, UBE2C, UBE2S,.
M prospg UP
GO:0006260~DNA replication
POLD3, SUPT16H, PCNA, MCM7, NASP, CHEK1, MCM4, MCM5, ...
GO:0005634~nucleus
SLBP, PCNA, MCM7, CHEK1, DAZL, TJP1, NASP, ID4, MCM3,..
Spg
Diff. Spg.E
STRA8
REC8
PIWIL1
DNMT1
SYCP3
ASB9
EGR4
ID4
SOHLH1
UTF1
SIX1
FGFR3
PIWIL4
Spg UP
GO:0005667~transcription factor complex
SMAD1, JUND, MSX2, LMO4, SCX, GPS2, SMAD9, SIX1,..
GO:0009952~anterior/posterior pattern specification
TBX1, HEY1, MSX2, HOXA7, ..
Diff.Spg.E UP
GO:0005739~mitochondrion
NDUFA13, ACADVL, COX7B, MRPS16, ACAA2, TUSC3, COX4I1,.
GO:0009060~aerobic respiration
NDUFA13, NDUFB7, MT-ND5, UQCRB, NDUFA11, NDUFB5, NDUFB11, NDUFA3, NDUFB2, ..
M prospg
Int. prospg
MAP4K4
MDK
ASB9
ID3
STMN1
POU5F1
NANOS3
NANOG
MKI67
M prospg UP
GO:0051301~cell division
CDCA5, KIF14, NCAPG, BUB1B, KIF11, SMC4, CKS1B, CCNB2, ..
GO:0001714~endodermal cell fate specification
SOX17, NANOG, POU5F1
Int. prospg UP
GO:0003712~transcription cofactor activity
BTG1, NRIP1, CAMTA1, PBX1, SAP30, PHF8
GO:0007283~spermatogenesis
TDRD1, BTG1, CCNI, CADM1, YBX3, CFAP97, SEPTIN7
Diff.Spg.E
Diff.Spg.L
MKI67
HORMAD1
UBE2C
TEX15
SCML1
MEIOC
SYCP3
ASB9
KIT
L1TD1
ETV7
PIWIL1
Diff.Spg.E UP
GO:0005737~cytoplasm
PIWIL1, APP, RPL5, RPL30, RPL3, RPL32, CCNI, CITED2, FTH1, ..
GO:0045727~positive regulation of translation
RPS4X, RPL5, EEF2, SOX4,..
Diff.Spg.L UP
GO:0051301~cell division
SYCP2, KIF14, BUB1B, KIF11, SMC4, CKS1B, CDC20, TUBA1C, ..
GO:0007059~chromosome segregation
TOP2A, CENPU, CDCA2, CENPW, SPAG5, HJURP, TTK, ESCO2, ..
Int prospg
T1 prpspg
MAGEC2
SIX1
MAGEB2
TCF3
DDX4
STMN1
MEG3
APOE
POU5F1
NANOS3
ASB9
Int. prospg UP
GO:0005829~cytosol
EIF4A2, APP, CYFIP1, EIF4A1, RPL5, TCL1A, DDX17, NANOS3,..
GO:0042981~regulation of apoptotic process
ANP32A, DAPK1, APEX1, RPS3, NME2, BAX, CALR, APOE, TPT1, ..
T1 prospg UP
GO:0007283~spermatogenesis
PIWIL2, BCAP31, NANOS2, DDX4, CCNI, CSNK2A2, TEX15, YBX3, ..
GO:0034587~piRNA metabolic process
TDRD1, DDX4, HENMT1, MOV10L1
Diff.Spg.L
Prelep. Spc
SYCP1
TEX101
MEIOB
SYCP3
HORMAD1
ELAVL2
CENPF
ITGA6
RHOXF1
KIT
UBE2C
Diff.Spg.L UP
GO:0009060~aerobic respiration
MT-ND6, NDUFB9, MT-ND4L, NDUFA13, NDUFB8, MT-ND4, ...
GO:0070469~respiratory chain
MT-ND6, NDUFB9, MT-ND4L, NDUFB10, NDUFA12, NDUFA1,..
Prelep.Spc UP
GO:0007130~synaptonemal complex assembly
SYCP2, SPO11, SYCP1, RAD21L1, TEX12, TEX15, C1ORF146, ...
GO:0007141~male meiosis I
SPO11, HSPA2, BRCA2, ING2, RAD51C, UBB, DMC1, DMRTC2, ..

**Figure S6**

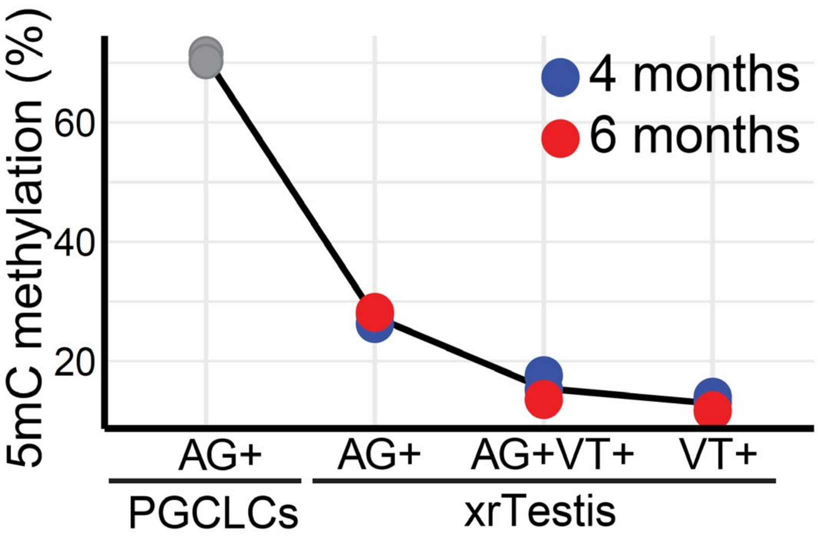

4 months
6 months
5mC methylation (%)
60
40
20
AG+
AG+
AG+VT+
VT+
PGCLCs
xrTestis

### Figure S7

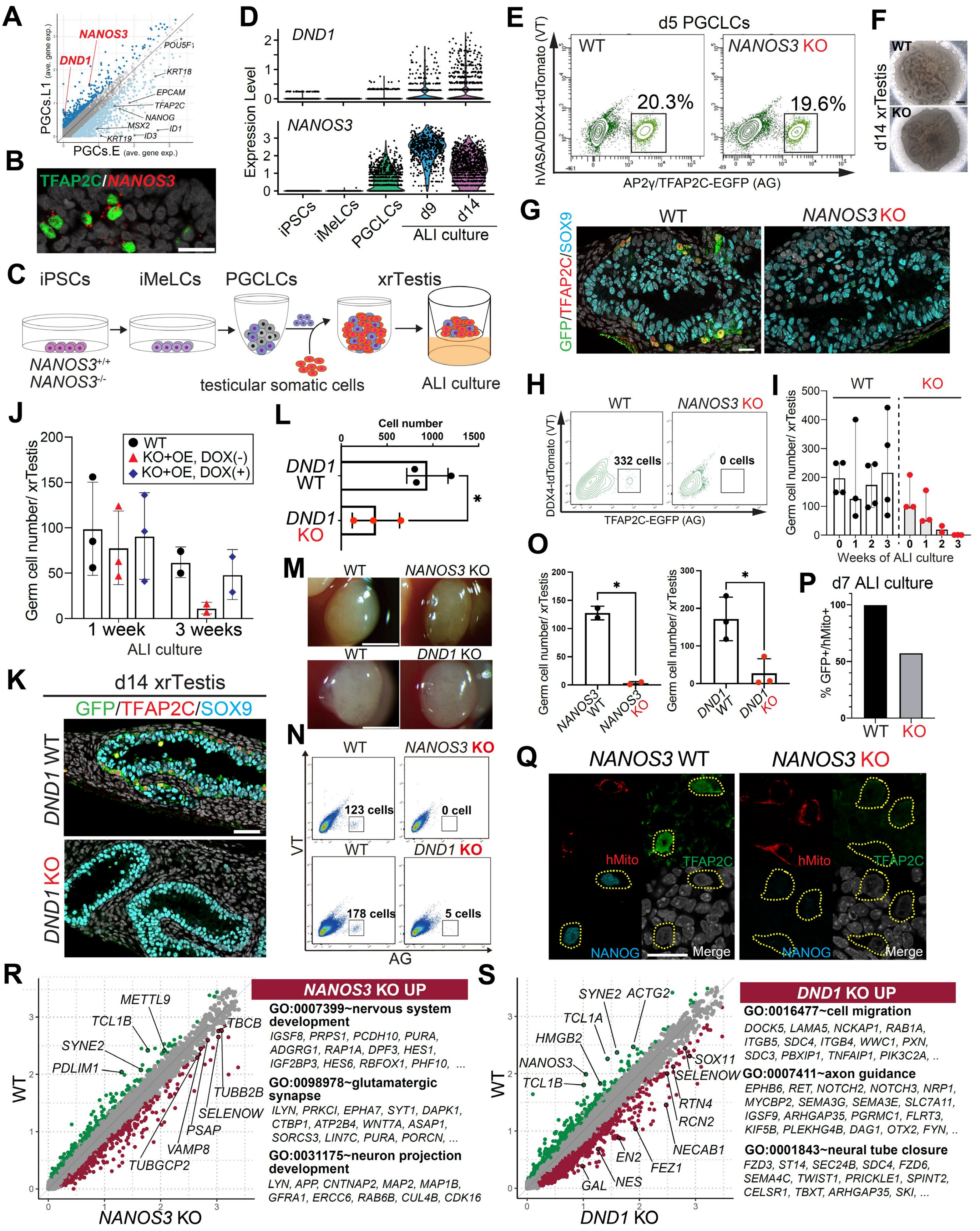
A
NANOS3
DND1
POU5F1
KRT18
EPCAM
TFAP2C
NANOG
MSX2
ID1
KRT19
ID3
PGCs.L1 (ave. gene exp.)
PGCs.E (ave. gene exp.)
B
TFAP2C/NANOS3
C
iPSCs
iMeLCs
PGCLCs
xrTestis
NANOS3+/+
NANOS3-/-
testicular somatic cells
ALI culture
D
DND1
NANOS3
Expression Level
iPSCs
iMeLCs
PGCLCs
d9
d14
ALI culture
E
d5 PGCLCs
WT
NANOS3 KO
20.3%
19.6%
hVASA/DDX4-tdTomato (VT)
AP2γ/TFAP2C-EGFP (AG)
F
WT
KO
d14 xrTestis
G
WT
NANOS3 KO
GFP/TFAP2C/SOX9
H
WT
NANOS3 KO
332 cells
0 cells
DDX4-tdTomato (VT)
TFAP2C-EGFP (AG)
I
WT
KO
Germ cell number/ xrTestis
Weeks of ALI culture
J
WT
KO+OE, DOX(-)
KO+OE, DOX(+)
Germ cell number/ xrTestis
1 week
3 weeks
ALI culture
L
Cell number
DND1 WT
DND1 KO
*
M
WT
NANOS3 KO
WT
DND1 KO
N
WT
NANOS3 KO
123 cells
0 cell
WT
DND1 KO
178 cells
5 cells
VT
AG
O
Germ cell number/ xrTestis
NANOS3 WT
NANOS3 KO
DND1 WT
DND1 KO
*
P
d7 ALI culture
% GFP+/hMito+
WT
KO
K
d14 xrTestis
GFP/TFAP2C/SOX9
DND1 WT
DND1 KO
Q
NANOS3 WT
NANOS3 KO
hMito
TFAP2C
NANOG
Merge
R
METTL9
TCL1B
SYNE2
PDLIM1
TBCB
TUBB2B
SELENOW
PSAP
VAMP8
TUBGCP2
WT
NANOS3 KO
NANOS3 KO UP
GO:0007399~nervous system development
IGSF8, PRPS1, PCDH10, PURA, ADGRG1, RAP1A, DPF3, HES1, IGF2BP3, HES6, RBFOX1, PHF10, ...
GO:0098978~glutamatergic synapse
ILYN, PRKCI, EPHA7, SYT1, DAPK1, CTBP1, ATP2B4, WNT7A, ASAP1, SORCS3, LIN7C, PURA, PORCN, ...
GO:0031175~neuron projection development
LYN, APP, CNTNAP2, MAP2, MAP1B, GFRA1, ERCC6, RAB6B, CUL4B, CDK16
S
SYNE2
ACTG2
TCL1A
HMGB2
NANOS3
TCL1B
SOX11
SELENOW
RTN4
RCN2
NECAB1
EN2
FEZ1
NES
GAL
WT
DND1 KO
DND1 KO UP
GO:0016477~cell migration
DOCK5, LAMA5, NCKAP1, RAB1A, ITGB5, SDC4, ITGB4, WWC1, PXN, SDC3, PBXIP1, TNFAIP1, PIK3C2A, ..
GO:0007411~axon guidance
EPHB6, RET, NOTCH2, NOTCH3, NRP1, MYCBP2, SEMA3G, SEMA3E, SLC7A11, IGSF9, ARHGAP35, PGRMC1, FLRT3, KIF5B, PLEKHG4B, DAG1, OTX2, FYN, ..
GO:0001843~neural tube closure
FZD3, ST14, SEC24B, SDC4, FZD6, SEMA4C, TWIST1, PRICKLE1, SPINT2, CELSR1, TBXT, ARHGAP35, SKI, ...

### Figure S8

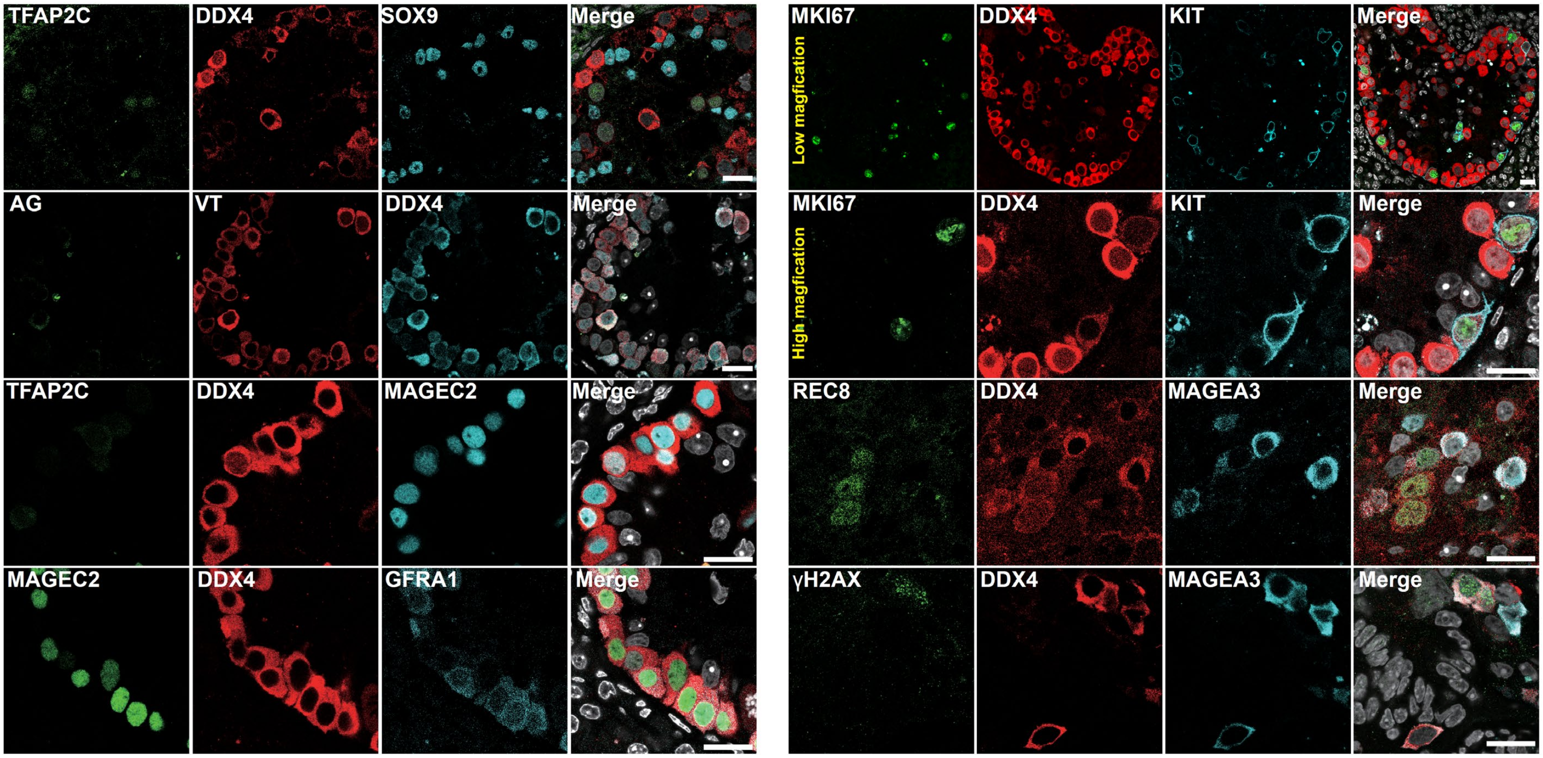
TFAP2C
DDX4
SOX9
Merge
AG
VT
DDX4
Merge
TFAP2C
DDX4
MAGEC2
Merge
MAGEC2
DDX4
GFRA1
Merge
MKI67
Low magfication
DDX4
KIT
Merge
MKI67
High magfication
DDX4
KIT
Merge
REC8
DDX4
MAGEA3
Merge
γH2AX
DDX4
MAGEA3
Merge

### Figure S9

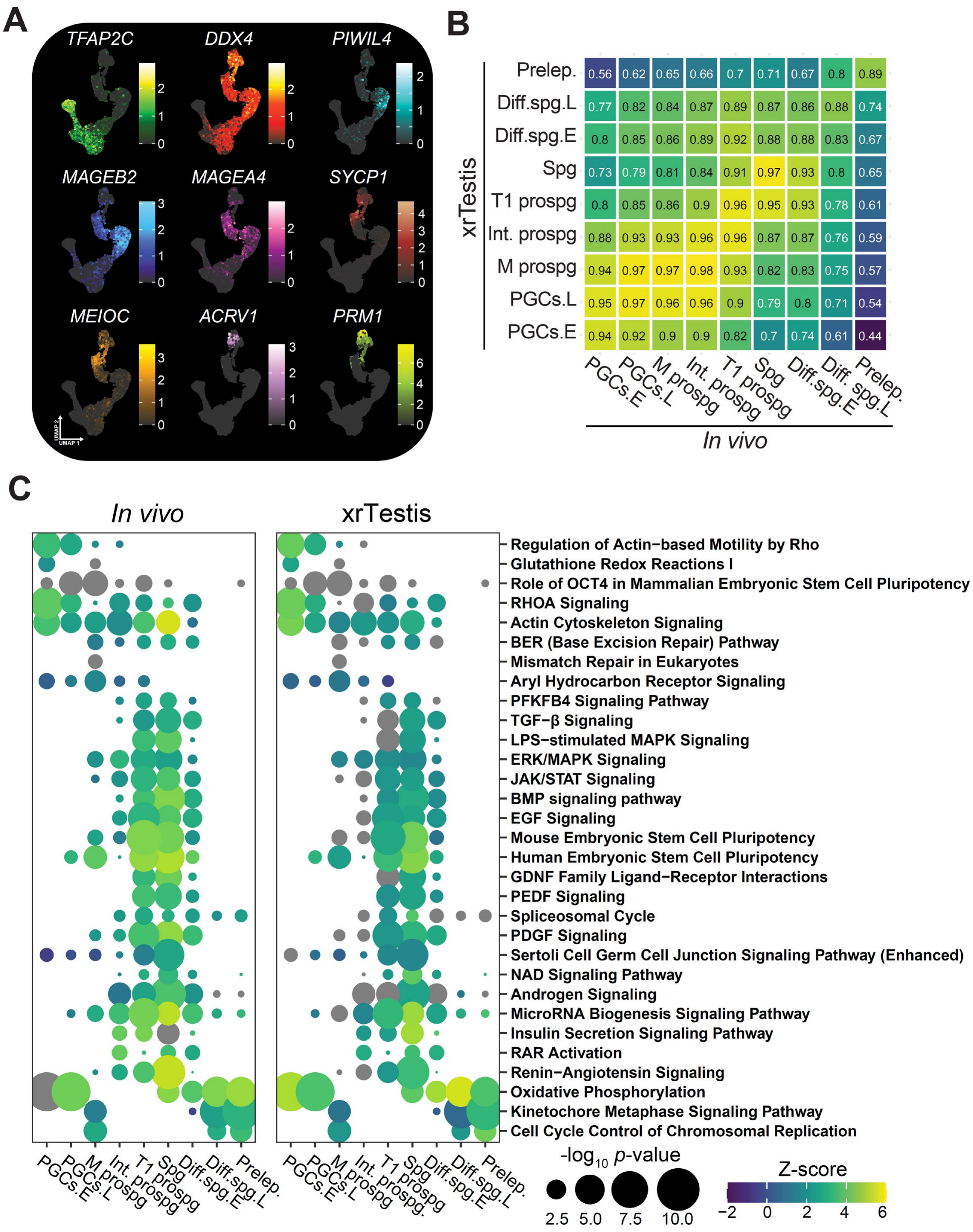
A
TFAP2C
DDX4
PIWIL4
MAGEB2
MAGEA4
SYCP1
MEIOC
ACRV1
PRM1
UMAP 2
UMAP 1
B
xrTestis
Prelep. 0.56 0.62 0.65 0.66 0.7 0.71 0.67 0.8 0.89
Diff.spg.L 0.77 0.82 0.84 0.87 0.89 0.87 0.86 0.88 0.74
Diff.spg.E 0.8 0.85 0.86 0.89 0.92 0.88 0.88 0.83 0.67
Spg 0.73 0.79 0.81 0.84 0.91 0.97 0.93 0.8 0.65
T1 prospg 0.8 0.85 0.86 0.9 0.96 0.95 0.93 0.78 0.61
Int. prospg 0.88 0.93 0.93 0.96 0.96 0.87 0.87 0.76 0.59
M prospg 0.94 0.97 0.97 0.98 0.93 0.82 0.83 0.75 0.57
PGCs.L 0.95 0.97 0.96 0.96 0.9 0.79 0.8 0.71 0.54
PGCs.E 0.94 0.92 0.9 0.9 0.82 0.7 0.74 0.61 0.44
PGCs.E PGCs.L M prospg Int. prospg T1 prospg Spg Diff.spg.E Diff. spg.L Prelep.
In vivo
C
In vivo
xrTestis
Regulation of Actin−based Motility by Rho
Glutathione Redox Reactions I
Role of OCT4 in Mammalian Embryonic Stem Cell Pluripotency
RHOA Signaling
Actin Cytoskeleton Signaling
BER (Base Excision Repair) Pathway
Mismatch Repair in Eukaryotes
Aryl Hydrocarbon Receptor Signaling
PFKFB4 Signaling Pathway
TGF−β Signaling
LPS−stimulated MAPK Signaling
ERK/MAPK Signaling
JAK/STAT Signaling
BMP signaling pathway
EGF Signaling
Mouse Embryonic Stem Cell Pluripotency
Human Embryonic Stem Cell Pluripotency
GDNF Family Ligand−Receptor Interactions
PEDF Signaling
Spliceosomal Cycle
PDGF Signaling
Sertoli Cell Germ Cell Junction Signaling Pathway (Enhanced)
NAD Signaling Pathway
Androgen Signaling
MicroRNA Biogenesis Signaling Pathway
Insulin Secretion Signaling Pathway
RAR Activation
Renin−Angiotensin Signaling
Oxidative Phosphorylation
Kinetochore Metaphase Signaling Pathway
Cell Cycle Control of Chromosomal Replication
PGCs.E PGCs.L M prospg Int. prospg T1 prospg Spg Diff.spg.E Diff.spg.L Prelep.
PGCs.E PGCs.L M prospg Int. prospg. T1 prospg Spg Diff.spg.E Diff.spg.L Prelep.
-log10 p-value
2.5 5.0 7.5 10.0
Z-score
−2 0 2 4 6

### Figure S10

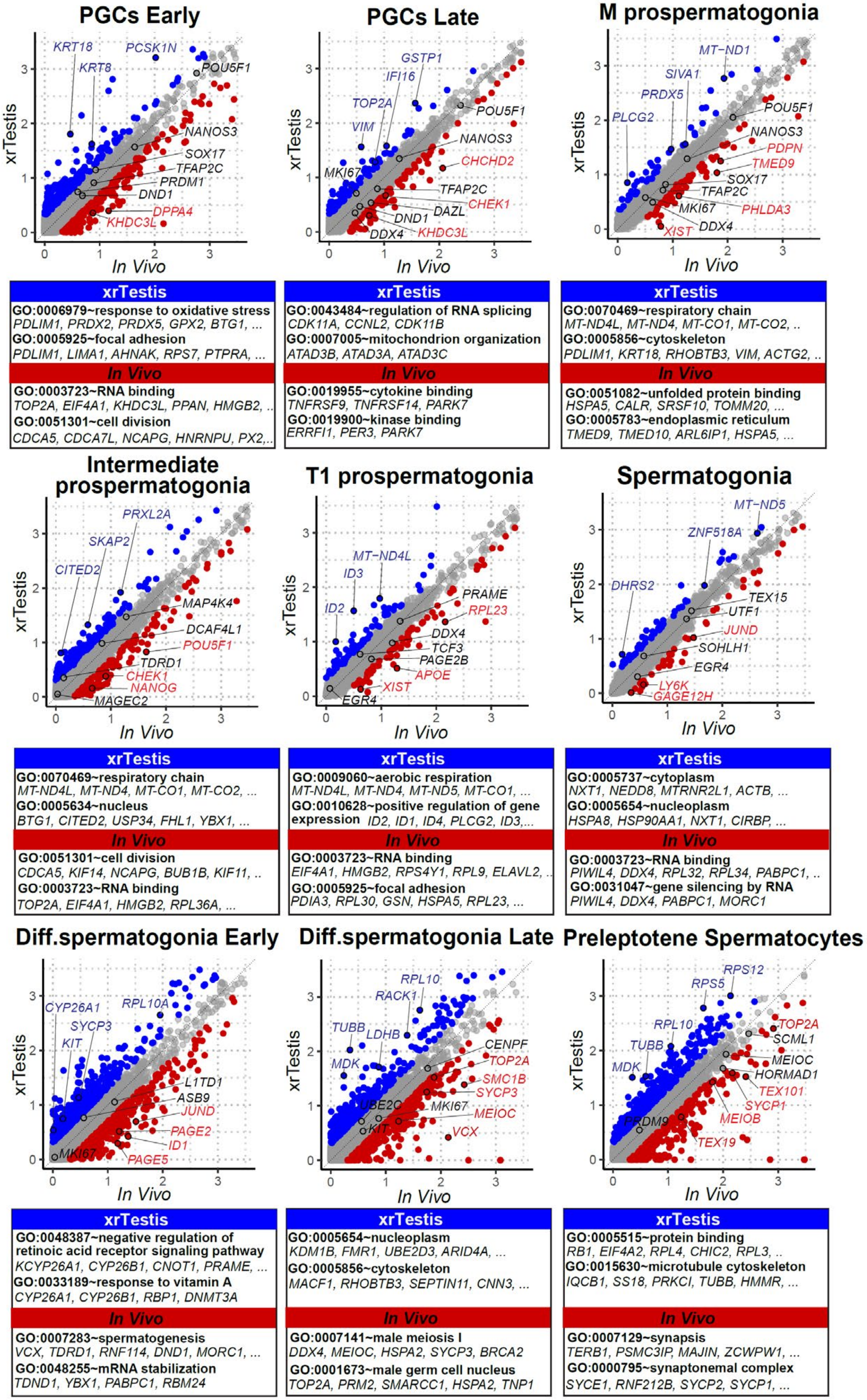

PGCs Early
KRT18
PCSK1N
KRT8
POU5F1
NANOS3
SOX17
TFAP2C
PRDM1
DND1
DPPA4
KHDC3L
xrTestis
In Vivo
PGCs Late
GSTP1
IFI16
TOP2A
VIM
POU5F1
NANOS3
CHCHD2
MKI67
TFAP2C
CHEK1
DAZL
DND1
DDX4
KHDC3L
M prospermatogonia
MT−ND1
SIVA1
PRDX5
PLCG2
POU5F1
NANOS3
PDPN
TMED9
SOX17
TFAP2C
MKI67
PHLDA3
DDX4
XIST
xrTestis
GO:0006979~response to oxidative stress
PDLIM1, PRDX2, PRDX5, GPX2, BTG1, ...
GO:0005925~focal adhesion
PDLIM1, LIMA1, AHNAK, RPS7, PTPRA, ...
In Vivo
GO:0003723~RNA binding
TOP2A, EIF4A1, KHDC3L, PPAN, HMGB2, ..
GO:0051301~cell division
CDCA5, CDCA7L, NCAPG, HNRNPU, PX2,..
xrTestis
GO:0043484~regulation of RNA splicing
CDK11A, CCNL2, CDK11B
GO:0007005~mitochondrion organization
ATAD3B, ATAD3A, ATAD3C
In Vivo
GO:0019955~cytokine binding
TNFRSF9, TNFRSF14, PARK7
GO:0019900~kinase binding
ERRFI1, PER3, PARK7
xrTestis
GO:0070469~respiratory chain
MT-ND4L, MT-ND4, MT-CO1, MT-CO2, ..
GO:0005856~cytoskeleton
PDLIM1, KRT18, RHOBTB3, VIM, ACTG2, ..
In Vivo
GO:0051082~unfolded protein binding
HSPA5, CALR, SRSF10, TOMM20, ...
GO:0005783~endoplasmic reticulum
TMED9, TMED10, ARL6IP1, HSPA5, ...
Intermediate prospermatogonia
PRXL2A
SKAP2
CITED2
MAP4K4
DCAF4L1
POU5F1
TDRD1
CHEK1
NANOG
MAGEC2
T1 prospermatogonia
MT−ND4L
ID3
ID2
PRAME
RPL23
DDX4
TCF3
PAGE2B
APOE
XIST
EGR4
Spermatogonia
MT−ND5
ZNF518A
DHRS2
TEX15
UTF1
JUND
SOHLH1
EGR4
LY6K
GAGE12H
xrTestis
GO:0070469~respiratory chain
MT-ND4L, MT-ND4, MT-CO1, MT-CO2, ...
GO:0005634~nucleus
BTG1, CITED2, USP34, FHL1, YBX1, ...
In Vivo
GO:0051301~cell division
CDCA5, KIF14, NCAPG, BUB1B, KIF11, ..
GO:0003723~RNA binding
TOP2A, EIF4A1, HMGB2, RPL36A, ...
xrTestis
GO:0009060~aerobic respiration
MT-ND4L, MT-ND4, MT-ND5, MT-CO1, ...
GO:0010628~positive regulation of gene expression ID2, ID1, ID4, PLCG2, ID3,...
In Vivo
GO:0003723~RNA binding
EIF4A1, HMGB2, RPS4Y1, RPL9, ELAVL2, ..
GO:0005925~focal adhesion
PDIA3, RPL30, GSN, HSPA5, RPL23, ...
xrTestis
GO:0005737~cytoplasm
NXT1, NEDD8, MTRNR2L1, ACTB, ...
GO:0005654~nucleoplasm
HSPA8, HSP90AA1, NXT1, CIRBP, ...
In Vivo
GO:0003723~RNA binding
PIWIL4, DDX4, RPL32, RPL34, PABPC1, ..
GO:0031047~gene silencing by RNA
PIWIL4, DDX4, PABPC1, MORC1
Diff.spermatogonia Early
CYP26A1
RPL10A
SYCP3
KIT
L1TD1
ASB9
JUND
PAGE2
ID1
MKI67
PAGE5
Diff.spermatogonia Late
RPL10
RACK1
TUBB
LDHB
MDK
CENPF
TOP2A
SMC1B
SYCP3
UBE2C
MKI67
MEIOC
KIT
VCX
Preleptotene Spermatocytes
RPS12
RPS5
TOP2A
SCML1
RPL10
TUBB
MEIOC
MDK
HORMAD1
TEX101
SYCP1
MEIOB
PRDM9
TEX19
xrTestis
GO:0048387~negative regulation of retinoic acid receptor signaling pathway
KCYP26A1, CYP26B1, CNOT1, PRAME, ...
GO:0033189~response to vitamin A
CYP26A1, CYP26B1, RBP1, DNMT3A
In Vivo
GO:0007283~spermatogenesis
VCX, TDRD1, RNF114, DND1, MORC1, ...
GO:0048255~mRNA stabilization
TDND1, YBX1, PABPC1, RBM24
xrTestis
GO:0005654~nucleoplasm
KDM1B, FMR1, UBE2D3, ARID4A, ...
GO:0005856~cytoskeleton
MACF1, RHOBTB3, SEPTIN11, CNN3, ...
In Vivo
GO:0007141~male meiosis I
DDX4, MEIOC, HSPA2, SYCP3, BRCA2
GO:0001673~male germ cell nucleus
TOP2A, PRM2, SMARCC1, HSPA2, TNP1
xrTestis
GO:0005515~protein binding
RB1, EIF4A2, RPL4, CHIC2, RPL3, ..
GO:0015630~microtubule cytoskeleton
IQCB1, SS18, PRKCI, TUBB, HMMR, ...
In Vivo
GO:0007129~synapsis
TERB1, PSMC3IP, MAJIN, ZCWPW1, ...
GO:0000795~synaptonemal complex
SYCE1, RNF212B, SYCP2, SYCP1, ...

### Figure S11

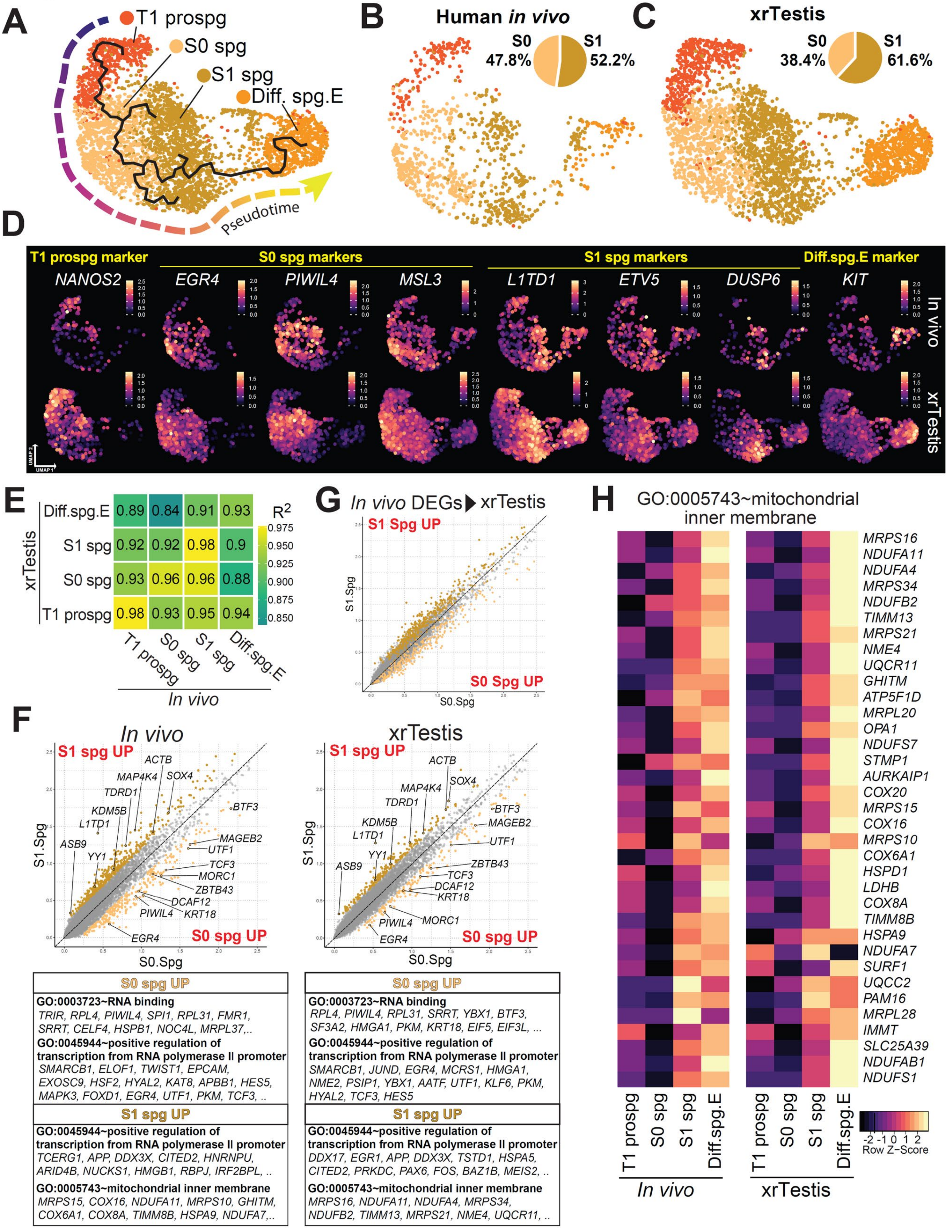
A
T1 prospg
S0 spg
S1 spg
Diff. spg.E
Pseudotime
B
Human in vivo
S0 47.8%
S1 52.2%
C
xrTestis
S0 38.4%
S1 61.6%
D
T1 prospg marker
S0 spg markers
S1 spg markers
Diff.spg.E marker
NANOS2
EGR4
PIWIL4
MSL3
L1TD1
ETV5
DUSP6
KIT
In vivo
xrTestis
UMAP 1
UMAP 2
E
xrTestis
Diff.spg.E 0.89 0.84 0.91 0.93
S1 spg 0.92 0.92 0.98 0.9
S0 spg 0.93 0.96 0.96 0.88
T1 prospg 0.98 0.93 0.95 0.94
T1 prospg
S0 spg
S1 spg
Diff.spg.E
In vivo
$R^2$
0.975
0.950
0.925
0.900
0.875
0.850
G
In vivo DEGs ▶ xrTestis
S1 Spg UP
S0 Spg UP
S1.Spg
S0.Spg
H
GO:0005743~mitochondrial inner membrane
MRPS16
NDUFA11
NDUFA4
MRPS34
NDUFB2
TIMM13
MRPS21
NME4
UQCR11
GHITM
ATP5F1D
MRPL20
OPA1
NDUFS7
STMP1
AURKAIP1
COX20
MRPS15
COX16
MRPS10
COX6A1
HSPD1
LDHB
COX8A
TIMM8B
HSPA9
NDUFA7
SURF1
UQCC2
PAM16
MRPL28
IMMT
SLC25A39
NDUFAB1
NDUFS1
T1 prospg
S0 spg
S1 spg
Diff.spg.E
In vivo
xrTestis
-2 -1 0 1 2
Row Z-Score
F
In vivo
S1 spg UP
S0 spg UP
ACTB
MAP4K4
SOX4
TDRD1
KDM5B
L1TD1
ASB9
YY1
BTF3
MAGEB2
UTF1
TCF3
MORC1
ZBTB43
DCAF12
PIWIL4
KRT18
EGR4
S1.Spg
S0.Spg
xrTestis
S1 spg UP
S0 spg UP
ACTB
SOX4
MAP4K4
TDRD1
KDM5B
L1TD1
YY1
ASB9
BTF3
MAGEB2
UTF1
ZBTB43
TCF3
DCAF12
KRT18
MORC1
PIWIL4
EGR4
S1.Spg
S0.Spg
S0 spg UP
GO:0003723~RNA binding
TRIR, RPL4, PIWIL4, SPI1, RPL31, FMR1, SRRT, CELF4, HSPB1, NOC4L, MRPL37,..
GO:0045944~positive regulation of transcription from RNA polymerase II promoter
SMARCB1, ELOF1, TWIST1, EPCAM, EXOSC9, HSF2, HYAL2, KAT8, APBB1, HES5, MAPK3, FOXD1, EGR4, UTF1, PKM, TCF3, ..
S1 spg UP
GO:0045944~positive regulation of transcription from RNA polymerase II promoter
TCERG1, APP, DDX3X, CITED2, HNRNPU, ARID4B, NUCKS1, HMGB1, RBPJ, IRF2BPL, ..
GO:0005743~mitochondrial inner membrane
MRPS15, COX16, NDUFA11, MRPS10, GHITM, COX6A1, COX8A, TIMM8B, HSPA9, NDUFA7,..
S0 spg UP
GO:0003723~RNA binding
RPL4, PIWIL4, RPL31, SRRT, YBX1, BTF3, SF3A2, HMGA1, PKM, KRT18, EIF5, EIF3L, ...
GO:0045944~positive regulation of transcription from RNA polymerase II promoter
SMARCB1, JUND, EGR4, MCRS1, HMGA1, NME2, PSIP1, YBX1, AATF, UTF1, KLF6, PKM, HYAL2, TCF3, HES5
S1 spg UP
GO:0045944~positive regulation of transcription from RNA polymerase II promoter
DDX17, EGR1, APP, DDX3X, TSTD1, HSPA5, CITED2, PRKDC, PAX6, FOS, BAZ1B, MEIS2, ..
GO:0005743~mitochondrial inner membrane
MRPS16, NDUFA11, NDUFA4, MRPS34, NDUFB2, TIMM13, MRPS21, NME4, UQCR11, ..

### Figure S12

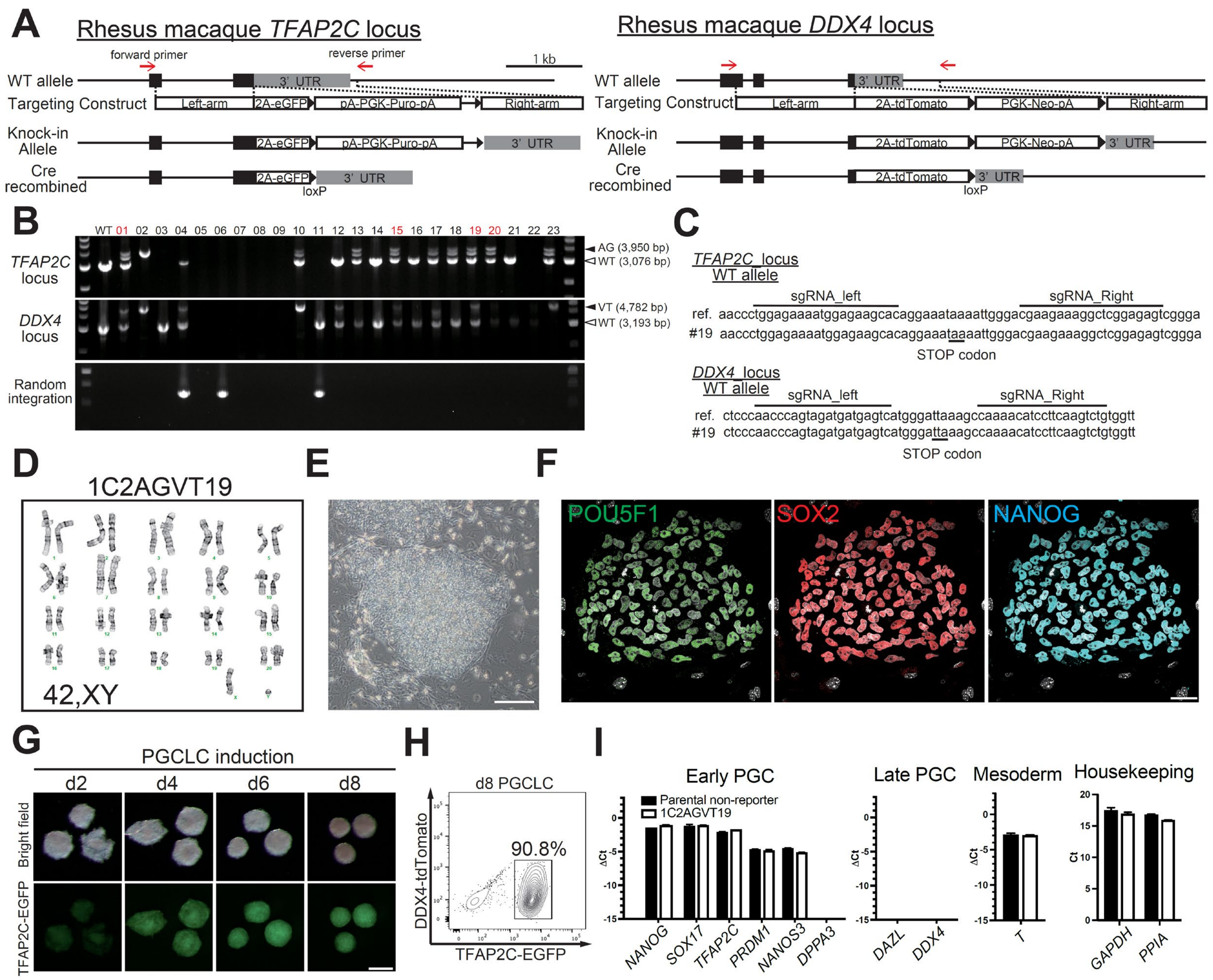

A
Rhesus macaque TFAP2C locus
forward primer
reverse primer
1 kb
WT allele
3' UTR
Targeting Construct
Left-arm
2A-eGFP
pA-PGK-Puro-pA
Right-arm
Knock-in Allele
Cre recombined
loxP
Rhesus macaque DDX4 locus
2A-tdTomato
PGK-Neo-pA
B
WT 01 02 03 04 05 06 07 08 09 10 11 12 13 14 15 16 17 18 19 20 21 22 23
TFAP2C locus
DDX4 locus
Random integration
AG (3,950 bp)
WT (3,076 bp)
VT (4,782 bp)
WT (3,193 bp)
C
TFAP2C locus
WT allele
sgRNA_left
sgRNA_Right
ref. aaccctggagaaaatggagaagcacaggaaataaaattgggacgaagaaaggctcggagagtcggga
#19 aaccctggagaaaatggagaagcacaggaaataaaattgggacgaagaaaggctcggagagtcggga
STOP codon
DDX4 locus
WT allele
ref. ctcccaacccagtagatgatgagtcatgggattaaagccaaaacatccttcaagtctgtggtt
#19 ctcccaacccagtagatgatgagtcatgggattaaagccaaaacatccttcaagtctgtggtt
STOP codon
D
1C2AGVT19
42,XY
E
F
POU5F1
SOX2
NANOG
G
PGCLC induction
d2
d4
d6
d8
Bright field
TFAP2C-EGFP
H
d8 PGCLC
90.8%
DDX4-tdTomato
TFAP2C-EGFP
I
Early PGC
Parental non-reporter
1C2AGVT19
NANOG
SOX17
TFAP2C
PRDM1
NANOS3
DPPA3
Late PGC
DAZL
DDX4
Mesoderm
T
Housekeeping
GAPDH
PPIA
ΔCt
Ct

### Figure S13

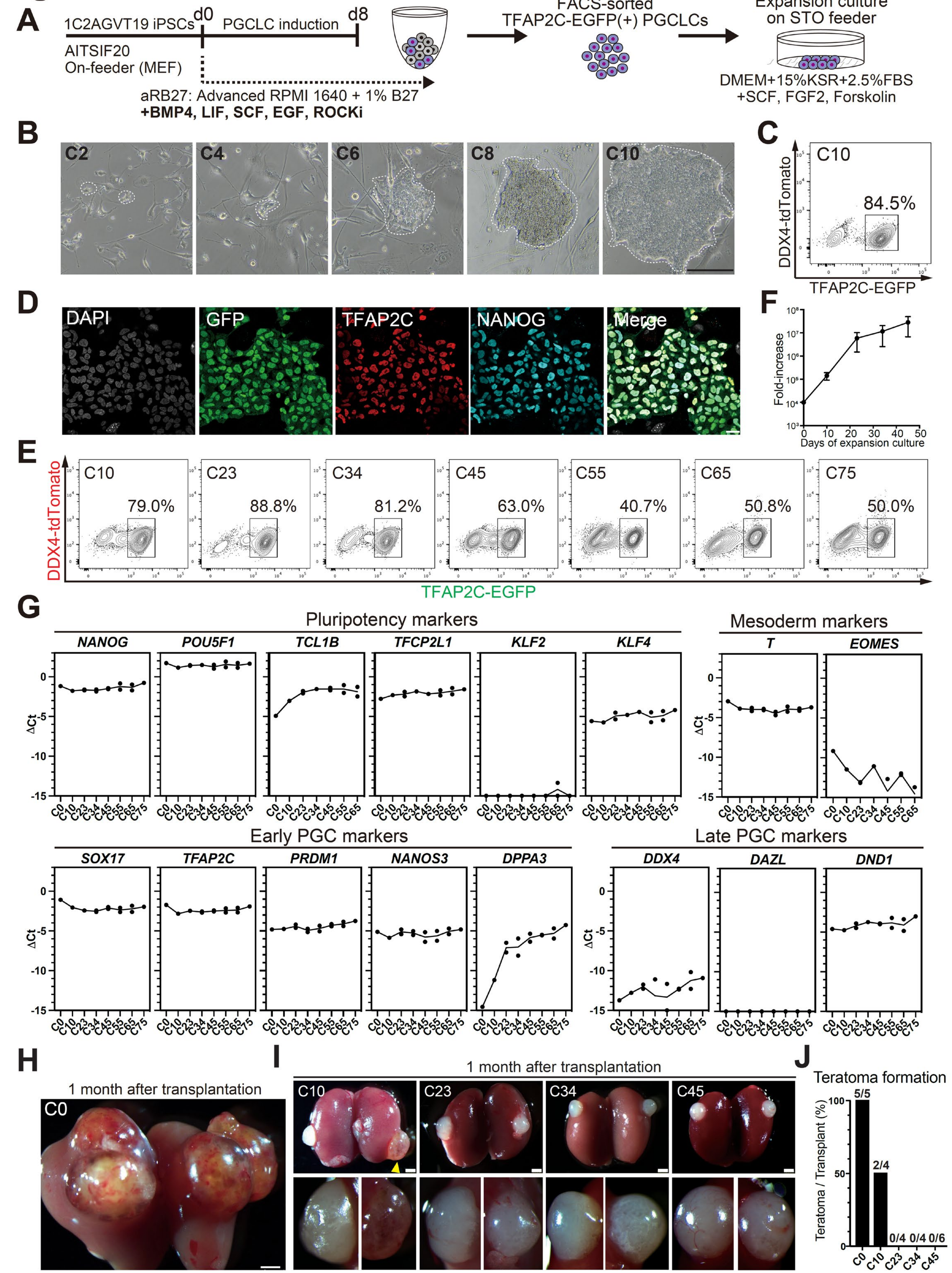

A
1C2AGVT19 iPSCs
d0
PGCLC induction
d8
AITSIF20
On-feeder (MEF)
aRB27: Advanced RPMI 1640 + 1% B27
+BMP4, LIF, SCF, EGF, ROCKi
FACS-sorted
TFAP2C-EGFP(+) PGCLCs
Expansion culture
on STO feeder
DMEM+15%KSR+2.5%FBS
+SCF, FGF2, Forskolin
B
C2
C4
C6
C8
C10
C
C10
84.5%
DDX4-tdTomato
TFAP2C-EGFP
D
DAPI
GFP
TFAP2C
NANOG
Merge
F
Fold-increase
Days of expansion culture
E
C10 79.0%
C23 88.8%
C34 81.2%
C45 63.0%
C55 40.7%
C65 50.8%
C75 50.0%
DDX4-tdTomato
TFAP2C-EGFP
G
Pluripotency markers
NANOG
POU5F1
TCL1B
TFCP2L1
KLF2
KLF4
Mesoderm markers
T
EOMES
Early PGC markers
SOX17
TFAP2C
PRDM1
NANOS3
DPPA3
Late PGC markers
DDX4
DAZL
DND1
ΔCt
H
1 month after transplantation
C0
I
1 month after transplantation
C10
C23
C34
C45
J
Teratoma formation
Teratoma / Transplant (%)
5/5
2/4
0/4
0/4
0/6
C0
C10
C23
C34
C45
